# The common evolution of fungicide persistence in cryptococcosis patients

**DOI:** 10.64898/2026.08.26.747313

**Authors:** Yuyan Xie, Weixin Ke, Xin Fan, Heping Xu, Wenzhao Wang, Zhen Zeng, Na Zhang, Haoran Ma, Zhongjie Tang, Huawei Zhu, Lei Jiang, Bo Zhu, Guohui Shi, Sihan Yang, Minling Yu, Feiyi Liu, Xixian Wang, Shaojie Li, Jian Xu, Koon Ho Wong, Xinping Xu, Cui Hua Liu, David S. Perlin, Yingchun Xu, Chong Li, Meng Xiao, Linqi Wang

**Author notes:** Correspondence (M.X.); (W.K.); (Y. Xu); (C.L.); (L.W.). These authors contributed equally (Yuyan Xie, Weixin Ke, Xin Fan, Heping Xu).

## Abstract

Fungicide persistence, the ability of dormant fungal cells to survive lethal drug exposure, undermines treatment efficacy, yet its clinical evolution remains largely unexplored. We assembled a nationwide collection of clinical *Cryptococcus neoformans* isolates from China and revealed extensive inter-strain variation in amphotericin B (AmB) persistence. This variation can arise from common evolutionary events in patients, generating previously undescribed high-persistence cryptococcal variants refractory to AmB clearance. Genome-wide fitness landscape analysis shows that persistence-associated mutations generally impose minimal fitness costs, facilitating persistence evolution even in fungistatic-resistant and hypervirulent genetic backgrounds. Machine learning identified deficient *ACO2* expression as a key predictor of high-persistence clinical isolates. *ACO2* deficiency reproduces the high-persistence phenotype by promoting POPC accumulation, which competes with AmB for its target. Moreover, the clinical-stage antifungal T-2307 effectively eliminates high-persistence strains across diverse genetic backgrounds. Collectively, our findings unveil common evolution of fungicide persistence in cryptococcosis patients, representing a previously overlooked clinical concern.

## INTRODUCTION

Invasive fungal pathogens represent a growing threat to human health, yet the repertoire of first-line antifungal agents, particularly those with fungicidal activity, remains severely limited.^1–6^ Among these, amphotericin B (AmB) exhibits potent fungicidal activity against most fungal pathogens and, as such, serves as a last-line therapeutic option for many life-threatening invasive fungal diseases, including cryptococcosis.^7–10^ Cryptococcosis is primarily caused by *Cryptococcus neoformans*,^11–15^ the top-ranking WHO fungal priority pathogen.^16–18^ Notably, classical AmB resistance, defined by elevated minimum inhibitory concentrations (MICs),^19, 20^ is exceedingly rare in clinical isolates of *C. neoformans* and other fungal pathogens,^21–26^ largely because resistance mutations incur fitness costs that restrict their emergence and colonization within the host.^27, 28^ Nevertheless, since the introduction of AmB into clinical practice in the 1950s,^7^ ineffective fungal clearance following standard AmB-based therapy has been frequently observed in patients with cryptococcosis.^29–32^ Given the extreme rarity of AmB resistance variants in clinical settings, alternative adaptive mechanisms beyond conventional resistance may underlie this phenomenon and could represent important factors underlying the difficulty in eliminating fungal cells in patients, thereby contributing to treatment failure and relapses.^20, 33–39^

One adaptive strategy that microbial pathogens use to counter microbicidal drugs is drug persistence, a phenomenon in which a subpopulation of metabolically dormant cells survives transient exposure to lethal drug concentrations without a change in MIC.^40–50^ In various fungal pathogens, including *C. neoformans*, drug persistence has been shown to compromise AmB efficacy *in vitro* and *in vivo*.^51–58^ Nevertheless, critical questions regarding its clinical relevance remain: Does fungicide persistence evolve during invasive cryptococcal infection to generate difficult-to-eliminate variants? And if so, what are the determinants and fitness costs that shape its *in vivo* trajectory? Addressing these questions may provide insight into the underlying reasons for, and potential strategies against, ineffective clearance by fungicide-based therapy, a challenge that has remained an important clinical concern since the introduction of AmB.

## RESULTS

### Development of a high-throughput assay to quantify AmB persistence in 1,025 clinical *C. neoformans* isolates from 67 hospitals in China

To systematically assess AmB persistence levels among clinical *C. neoformans* isolates, we established a large-scale clinical isolate repository by collecting 1,025 *C. neoformans* isolates from 67 hospitals across 25 provinces in China between 2010 and 2023 through the China Hospital Invasive Fungal Surveillance Net (CHIF-NET) program. These isolates were obtained from 942 patients with confirmed cryptococcosis, including 80 patients from whom two or more isolates were collected (Figures 1A and S1A). The vast majority of isolates were recovered from normally sterile sites,^59^ predominantly cerebrospinal fluid (CSF) (58.244%) and blood (30.341%), and were obtained via pure culture, minimizing the likelihood of contamination (Figure S1B and Table S1).^60^

**Figure 1.**
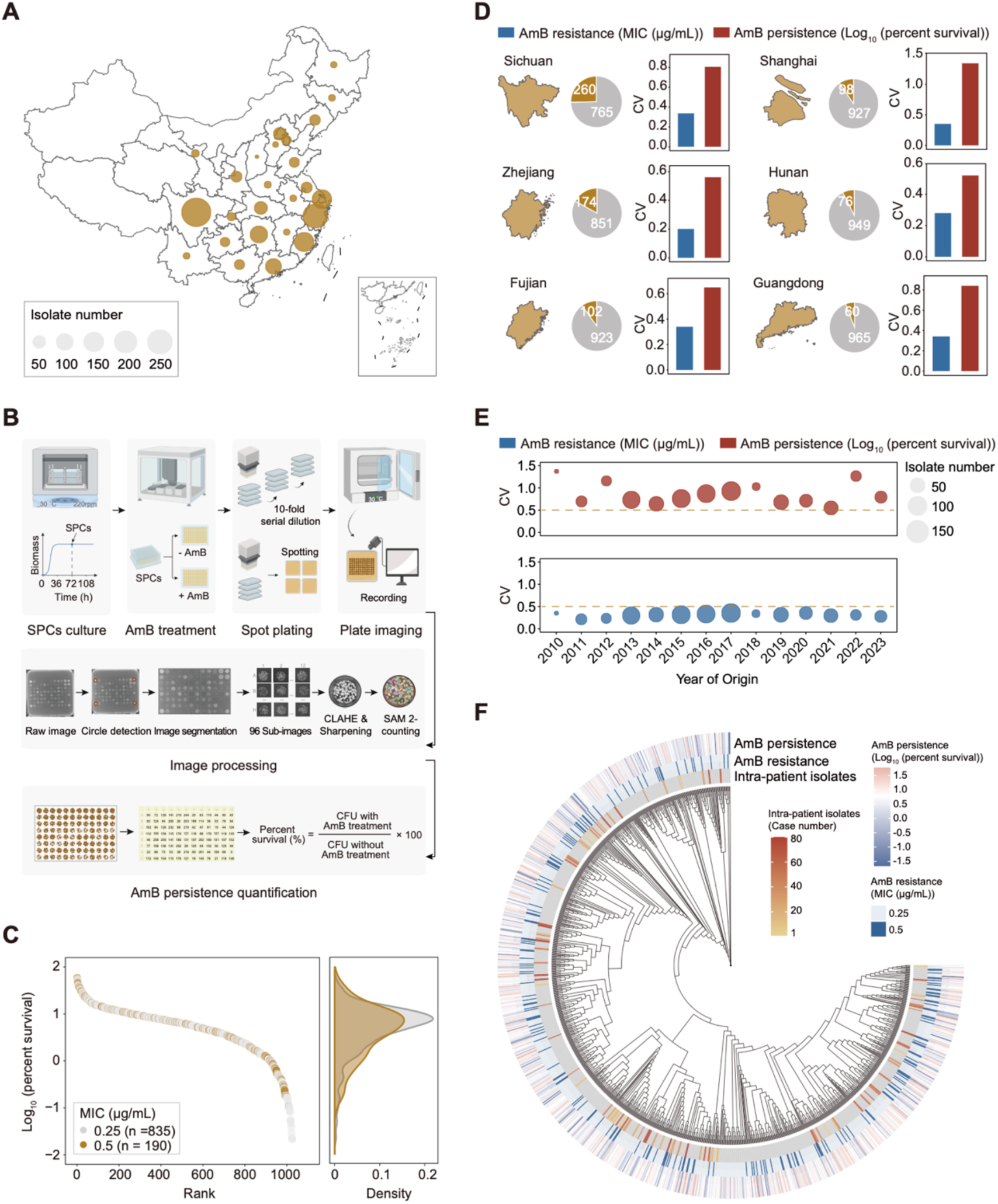
HiPERS enables high-throughput quantification of AmB persistence across a nationwide collection of clinical *C. neoformans* isolates. (A) Geographic distribution of the clinical *C. neoformans* isolates collected through the CHIF-NET program. (B) Experimental workflow of HiPERS. The figure was generated in BioRender. (C) AmB persistence levels of 1,025 clinical *C. neoformans* isolates, shown as log_10_ (percent survival) after 24-h-AmB treatment. Each point represents the mean value from two independent biological replicates. (D) AmB resistance and persistence profiles across the six provinces with more than 50 clinical isolates sampled. Pie charts indicate isolate numbers; bar plots show regional CVs. (E) AmB resistance and persistence profiles over the period from 2010 to 2023. (F) Whole-genome SNP-based phylogeny of 1,025 clinical *C. neoformans* isolates.

The previously reported persistence evaluation protocol typically involves culturing fungi to stationary-phase cells (SPCs) to obtain a relatively homogeneous population of dormant cells, followed by treatment with a high dose of AmB (exceeding the Clinical and Laboratory Standards Institute (CLSI) clinical breakpoint or epidemiological cutoff value (ECV) by more than 10-fold) for 24 h, and then quantifying persistence levels based on cell viability.^21, 54, 55, 57, 61–64^ However, this workflow is not suitable for large-scale strain collections due to throughput limitations. To address this, we developed a **H**igh-throughput **P**ersistence **E**valuation via Image **R**ecognition and **S**egmentation (HiPERS) pipeline (Figure 1B). This method employs 96-well plates combined with an automated platform for stationary-phase culture and drug incubation, after which cells from each well are then diluted and directly spotted onto drug-free agar plates (Figure 1B). A key challenge arises from the small spotting area derived from each well, where individual colonies tend to merge, making accurate quantification difficult with conventional colony-counting software. To overcome this challenge, we introduced image-recognition preprocessing algorithms including Contrast-Limited Adaptive Histogram Equalization (CLAHE) and sharpening, and applied the large-scale model Segment Anything Model 2 (SAM 2) for automated colony segmentation and fully automated counting (Figure 1B).^65–67^ By comparison, we demonstrated that our image analysis pipeline significantly outperforms other image-analysis approaches in quantification accuracy: the SAM 2-based pipeline achieved a mean absolute error (MAE) of 2.992 colonies per image with a coefficient of determination (R²) of 0.974 (Figure S1C).^68, 69^ Furthermore, a parallel comparison of 80 randomly selected strains showed that HiPERS yielded survival measurements highly consistent with the conventional method (Pearson’s *r* = 0.934, *p* < 0.001) (Figure S1D).

Using HiPERS, we profiled AmB persistence levels in all 1,025 clinical isolates. In parallel, we determined MIC values reflecting classical resistance using the standard broth microdilution method. Regarding resistance, all isolates exhibited MIC values ranging from 0.25 to 0.5 µg/mL, with none exceeding the CLSI ECV of 0.5 µg/mL (Table S1).^21^ This confirms that AmB-resistant *C. neoformans* isolates are extremely rare in clinical settings.^21–24^ In contrast to resistance, AmB persistence levels varied markedly among clinical isolates, with survival rates spanning over three orders of magnitude overall and differing even among isolates with identical MICs (Figure 1C). Moreover, AmB persistence showed consistently high coefficients of variation (CVs) (> 0.5) at both province and year levels (Figures 1D and 1E). Together, these findings indicate that, unlike resistance, AmB persistence levels vary greatly among clinical *C. neoformans* strains.

To investigate the relationship between persistence levels and phylogenetic background in these clinical isolates, we performed whole-genome sequencing (WGS) on all 1,025 isolates. For unbiased analysis of the genomic diversity of these isolates, we carried out *de novo* genome assembly.^70, 71^ The assemblies yielded an average contig number of 89, an average N50 of 266,733 bp, and an average genome size of 18.671 Mb (Table S1). Multilocus sequence typing (MLST) confirmed that ST5 (90.146%) and ST31 (6.927%) are the dominant sequence types in this isolate collection (Figure S2A and Table S1), consistent with previous epidemiological reports on clinical *C. neoformans* isolates in China.^72^ Subsequently, STRUCTURE clustering and principal component analysis (PCA) based on whole-genome single-nucleotide polymorphisms (SNPs) partitioned these isolates into five distinct genetic clusters (Figures S2B-S2D). However, no statistically significant differences in AmB persistence levels were observed among isolates from these distinct genetic clusters (Figure S2E). Consistently, mapping AmB persistence levels onto the whole-genome phylogeny revealed no apparent clustering of isolates with similar persistence levels (Figure 1F). These results indicate that AmB persistence levels are not significantly associated with phylogenetic background among clinical *C. neoformans* isolates.

### Evolution of AmB persistence occurs during cryptococcal infection

Phylogenetic analysis showed that most clinical isolates derived from the same patient clustered closely on the phylogenetic tree; only a few isolates from the same patient (involving 8 cases) were not positioned as close phylogenetic neighbors (Figure 1F). This suggests that, in the majority of cases, isolates from the same patient likely underwent clonal evolution from a single common ancestor.^73, 74^ Consistent with this hypothesis, whole-genome SNP analysis further revealed that isolates from the same patient were genetically closer to each other than to isolates from different patients (Figure 2A). Notably, most isolates from the same patient exhibited identical MICs but high variation in their AmB persistence levels (Figure 2B). These results raise the hypothesis that, unlike resistance (defined by changes in MIC), persistence could evolve within patients, giving rise to variants with divergent persistence capacities.

**Figure 2.**
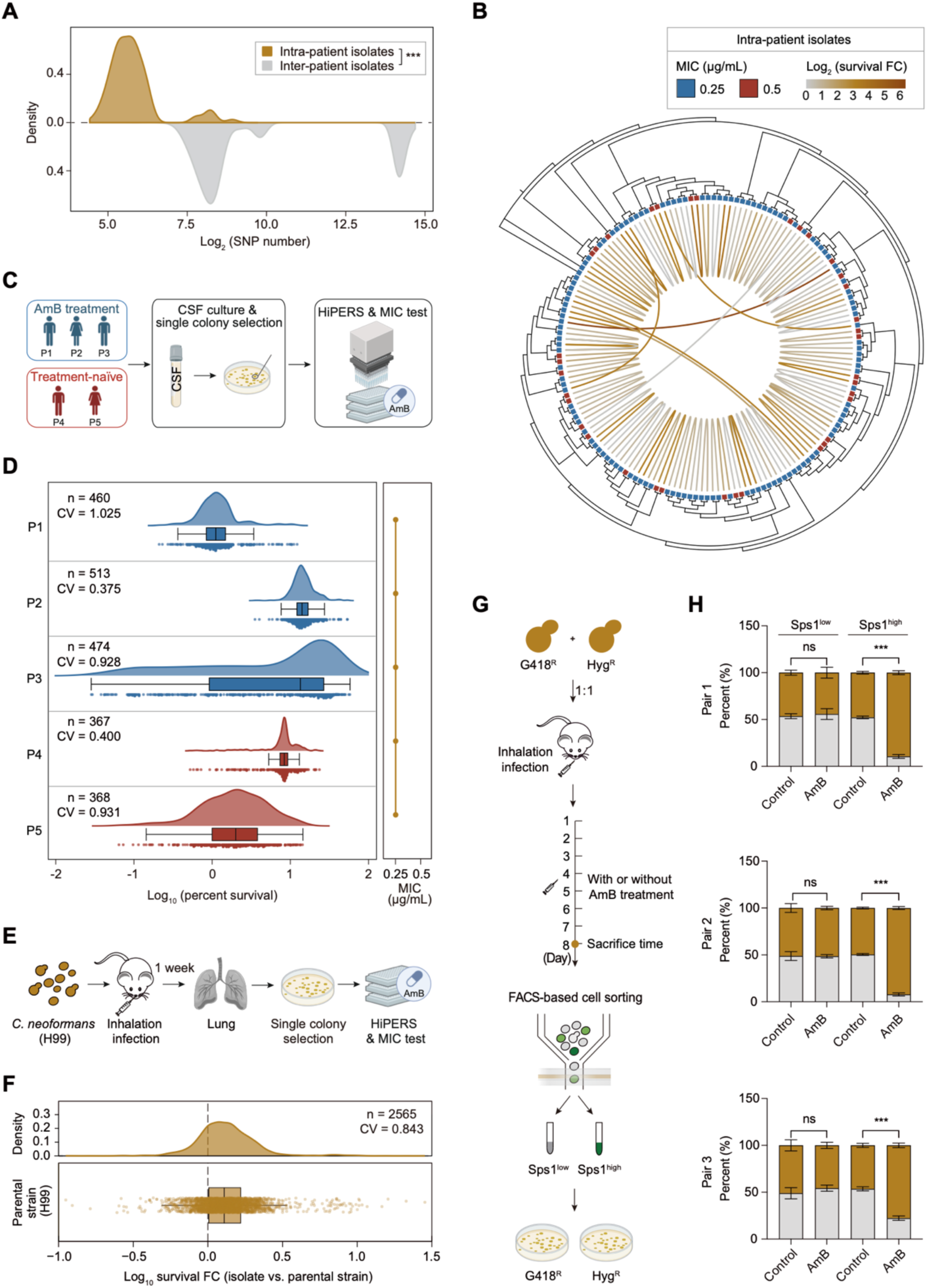
Within-host AmB persistence evolution promotes survival of dormant fungal cells under treatment. (A) Distribution of pairwise SNP distances between intra-patient isolates and inter-patient isolates, showing that isolates from the same patient are genetically closer than isolates from different patients. (B) Phylogenetic tree of 163 clinical *C. neoformans* isolates from 80 patients, with susceptibility to AmB (MICs) shown by blue/red boxes. Same-patient isolates are connected with lines whose color represent changes in AmB persistence. (C) Schematic diagram for profiling AmB susceptibility (MICs) and persistence across CSF-derived single-colony populations from five cryptococcosis patients (P1–P5). P1-P3 were sampled after AmB treatment, whereas P4 and P5 were treatment-naïve. The figure was generated in BioRender. (D) AmB susceptibility (MICs) and persistence profiles of CSF-derived single-colony populations from five cryptococcosis patients. CVs of AmB persistence within each patient-derived colony population are indicated. (E) Schematic diagram of *in vivo* evolution in a murine pulmonary cryptococcosis model. The figure was generated in BioRender. (F) AmB persistence profiles of lung-derived single-colony populations from infected mice. The MIC of the evolved isolates remained identical to that of the parental strain H99. (G) Schematic of the experimental workflow used to assess the *in vivo* killing efficacy of AmB against paired intra-patient isolates. (H) Relative proportions of paired intra-patient isolates in the cell populations with low Sps1 expression and high Sps1 expression after 7 days in mice with or without AmB treatment. Data shown as mean ± SD (n = 3). ns, not significant; \*\*\**p* < 0.001.

To test this hypothesis, we prospectively collected CSF samples from five patients with confirmed cryptococcosis (Figure 2C). These patients were from two independent medical centers and had no known epidemiological links or familial relationships. Among these patients, CSF samples from three patients were collected after approximately one week of AmB induction therapy, while the remaining two samples were obtained from treatment-naïve patients. For each CSF sample, we cultured the specimens on YPD agar to recover fungal cells and randomly selected hundreds of single colonies, the persistence characteristics of which were then systematically quantified using the HiPERS strategy (Figure 2C). The results demonstrated that AmB persistence varied among isolates from the same patient, with differences spanning one to three orders of magnitude, whereas their AmB susceptibility (as measured by MIC) was identical within each patient (Figure 2D). Importantly, such persistence variation was also observed in isolates from treatment-naïve patients who had not received AmB therapy (Figure 2D). These findings indicate that *C. neoformans* persistence commonly evolves within patients, and this evolution does not strictly depend on therapeutic drug selection pressure.

To further corroborate this hypothesis, we performed *in vivo* experimental evolution using a mouse model of cryptococcosis.^75–77^ Mice were infected with the reference strain H99 via intranasal inhalation. At 7 days post-infection (dpi), we harvested and homogenized lung tissues, plated the homogenates on YPD agar to recover fungal cells, and systematically quantified the persistence phenotypes of hundreds of randomly selected single colonies using the HiPERS strategy (Figure 2E). The data showed that AmB persistence can evolve during cryptococcal infection and, importantly, that some isolates exhibited significantly higher persistence levels than the parental strain, while their MICs remained comparable (Figure 2F).

To determine whether the differences in persistence capacities among within-host isolates directly correspond to the *in vivo* eradication efficiency of their dormant cells by AmB, we used a previously established dormancy reporter system (P*_SPS1_-GFP::SPS1*) based on the *C. neoformans* dormancy-specific protein Sps1.^57^ We introduced this system into three matched pairs of clinical isolates, each pair derived from a single patient and consisting of two isolates with markedly different persistence levels. To enable strain tracking during the murine co-infection, the two isolates in each pair were labeled with distinct selectable markers conferring resistance to geneticin (G418) or hygromycin B (Hyg). For each pair, the two reporter strains were then mixed at a 1:1 ratio and used to co-infect mice as previously mentioned (Figure 2G).^57^ After 7 days of AmB treatment, we observed significantly greater abundance of the strain with higher persistence capability than the low-persistence strain within the Sps1^high^ dormant cell subpopulation in all three isolate pairs; by contrast, no significant difference was observed in the metabolically active subpopulation (characterized by low Sps1 expression) (Figure 2H). Importantly, in untreated control mice, the two isolates showed no significant differences in relative abundance in either the Sps1^high^ or Sps1^low^ subpopulation (Figure 2H), indicating that their divergent persistence phenotypes were not accompanied by a detectable difference in *in vivo* pathogenicity fitness. These findings support the notion that the persistence capacity of within-patient variants is highly related to the *in vivo* elimination efficiency of their dormant cells by AmB.

### Functional genomics reveals the fitness landscape of AmB persistence-associated mutations across diverse growth conditions

Previous studies have shown that AmB resistance mutations impose a high fitness cost in *C. neoformans*, profoundly compromising the *in vivo* survival and colonization of resistant strains.^27, 78^ In contrast, AmB persistence evolves readily during infection (Figure 2F), suggesting that the determinants and associated *in vivo* fitness costs of persistence may fundamentally differ from those of resistance.

To test this hypothesis, we performed a genome-wide comparison of the fitness costs imposed by persistence-associated versus resistance-associated mutations. To this end, we leveraged a genome-scale deletion mutant library generated by Boucher *et al*.^79^ This library relies on DNA barcode technology, which previously enabled the systematic profiling of each mutant’s competitive growth, quantified as relative fitness scores via barcode sequencing, across 141 *in vitro* conditions including exposure to AmB and fungistatic agents, and one *in vivo* mouse lung infection condition.^79^ This comprehensive dataset defines the phenotypic landscape of *C. neoformans*, providing a resource for assessing the fitness costs associated with each gene deletion under these conditions (Figure 3A).

**Figure 3.**
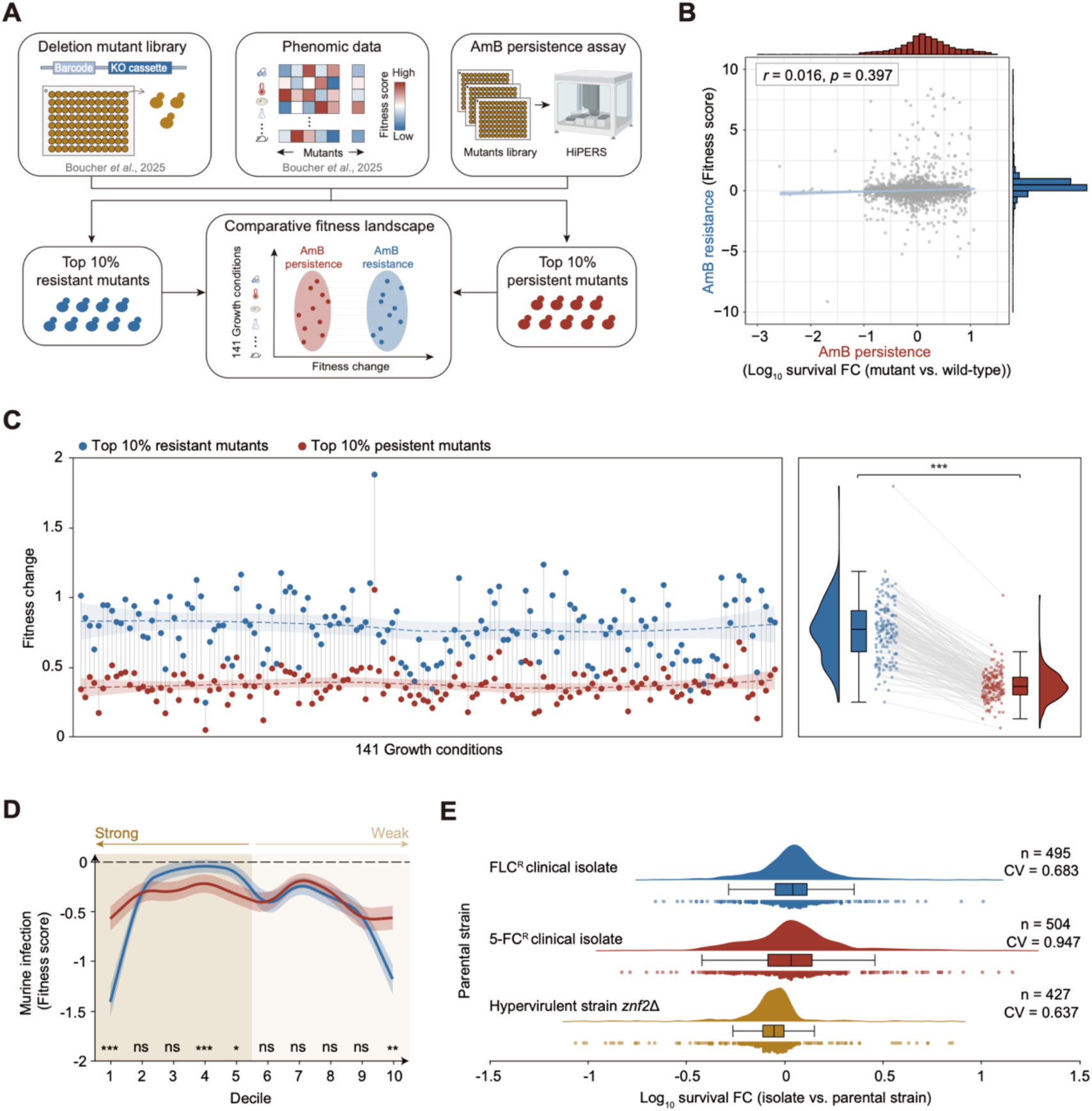
AmB persistence-associated mutations impose low fitness costs. (A) Schematic of the strategy used to compare the fitness landscapes of AmB persistence-associated and resistance-associated mutants. (B) Correlation between AmB persistence and AmB resistance across deletion mutants. AmB persistence is shown as the mean log_10_ (survival FC) from two independent biological replicates, whereas resistance is represented by the previously reported fitness score under AmB exposure. Pearson’s *r* and *p* value are indicated. (C) Fitness profiles of the top 10% AmB resistant and persistent mutants across 141 growth conditions. (D) Fitness scores in murine lung infection across deciles ranked by AmB resistance or AmB persistence. (E) *In vivo* evolution of AmB persistence in genetic backgrounds with clinically relevant traits, including fungistatic resistance or hypervirulence. ns, not significant; \**p* < 0.05; \*\**p* < 0.01; \*\*\**p* < 0.001.

Using the HiPERS strategy, we evaluated each mutant for its AmB persistence levels (Figure 3A). No significant correlation was observed between the AmB persistence levels and the previously reported AmB-exposed growth fitness-based resistance levels of these mutants (Pearson’s *r* = 0.016, *p* = 0.397) (Figure 3B),^79^ indicating that AmB persistence and resistance are governed by largely distinct genetic determinants.

We then compared the fitness landscapes of the top 10% of mutants ranked by AmB resistance (using the phenomic data generated by Boucher *et al*.) and the top 10% of mutants ranked by AmB persistence (as measured by HiPERS) (Table S2).^79^ The analysis revealed that, across all conditions, the top 10% resistant mutants exhibited significantly greater average fitness alterations than the top 10% persistent mutants (Figure 3C). Notably, when focusing specifically on the fitness cost in the murine lung infection, both the most resistant (top 10%) and the most susceptible (bottom 10%) mutants showed marked *in vivo* fitness defects (Figure 3D), consistent with the aforementioned findings regarding the rarity of both resistant and hyper-susceptible clinical isolates (Figure 1C). In comparison, both the top 10% and bottom 10% persistent mutants exhibited relatively minimal fitness costs (Figure 3D). These results indicate that, compared to resistance, AmB persistence-associated mutations typically impose only modest fitness burdens *in vivo*, which likely explains why the evolution of persistence occurs readily during infection.

Moreover, similar to murine lung infection, AmB persistence-associated mutations exerted a minimal impact on susceptibility to fungistatic agents such as fluconazole (FLC) and 5-flucytosine (5-FC) (Figures S3A and S3B). We therefore hypothesized that persistence evolution could occur in genetic backgrounds already exhibiting hypervirulence or resistance to fungistatic drugs, potentially generating strains with combined traits (e.g., fungistatic-resistant/AmB persistent or hypervirulent/AmB persistent). To test this, we selected a FLC-resistant (FLC^R^) clinical isolate (MIC = 32 µg/mL), a 5-FC-resistant (5-FC^R^) clinical isolate (MIC > 128 µg/mL), and a previously reported hypervirulent strain (*znf2*Δ in the H99 background),^80^ and established murine pulmonary infection models for each as previously mentioned (Figure 2E). One week post-infection, we recovered fungal isolates (more than 400 colonies per *in vivo* evolution experiment) from the lungs and quantified their persistence phenotypes using HiPERS. The results confirmed that both fungistatic-resistant and hypervirulent strains readily evolved *in vivo*, generating variants with significantly higher persistence levels than the parental strains (Figure 3E). Crucially, these evolved variants fully retained the fungistatic resistance or hypervirulence of their respective parental strains, as confirmed by susceptibility testing and mouse experiments (Figures S3C-S3E). These findings demonstrate that AmB persistence can evolve in genetic backgrounds already characterized by fungistatic resistance or hypervirulence, generating previously underappreciated variants that combine elevated persistence with other traits of clinical concern.

### Machine learning identifies low *ACO2* expression as a key predictor of high-persistence clinical isolates in *C. neoformans*

We next sought to identify the determinants underlying the high-persistence traits in clinical isolates. To address this question, we adopted a thresholding strategy inspired by the ECV, in which a cutoff defines the upper boundary of the wild-type phenotypic distribution and is typically set to include at least 95% of the wild-type population.^81^ Clinical isolates above this threshold, representing the top 5% with the highest survival rates, were classified as high-persistence (HP) strains, whereas the remaining isolates were classified as non-high-persistence (NHP) strains. We first examined the phylogenetic distribution of HP isolates and found that, instead of clustering within any specific clade, they were widely scattered across distinct sublineages (Figure 1F). Furthermore, a prediction model for HP isolates based on genome-wide variants (including both SNPs and insertions/deletions (InDels)) yielded modest performance (the area under the receiver operating characteristic curve (ROC-AUC) = 0.446 for Random Forest; ROC-AUC = 0.476 for XGBoost) (Figures S4A and S4B). These results indicate that clinical HP traits may have a complex genetic basis.

Recent studies have shown that integrating multimodal data with machine learning enables effective interpretation of complex biological traits.^82–84^ Accordingly, we selected 87 representative isolates from the 1,025 clinical isolates to generate multimodal datasets comprising genomic profiles, transcriptomic profiles, Raman spectroscopic fingerprints, and mass spectrometry-based features from their SPCs (Figure 4A and Table S3). Bootstrap resampling analysis showed that the median AmB persistence level of the selected 87 strains falls within the 95% resampling interval of random 87-strain subsets, supporting the representativeness of the selected strains (Figure 4B). Using Random Forest and XGBoost algorithms, we evaluated the predictive performance of single modalities and their pairwise fusions for the HP phenotype (Figure 4A).^85, 86^ Among these modalities, the transcriptomic expression profiles performed excellently, both alone and in combination with other modalities (Figure 4C).

**Figure 4.**
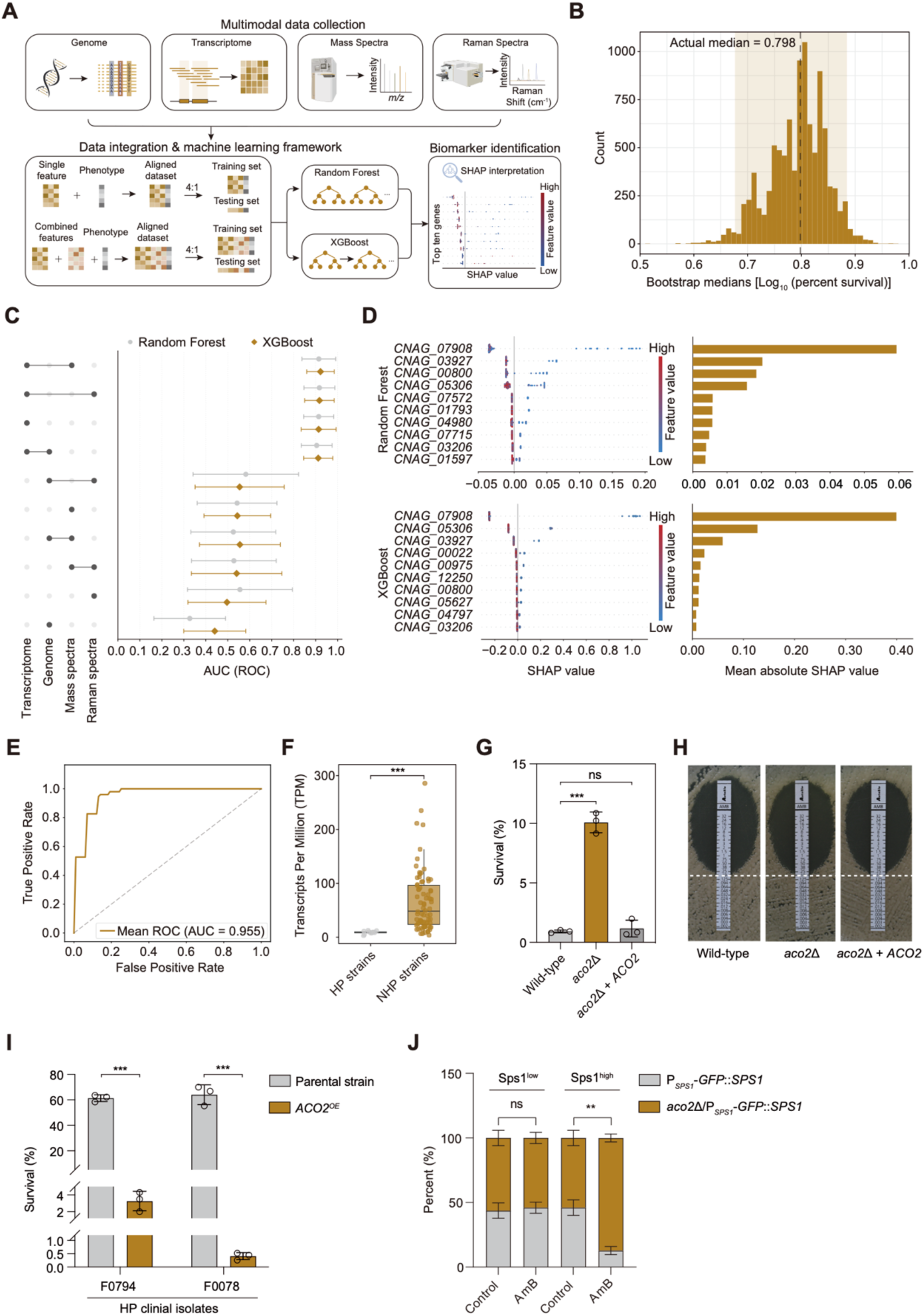
Multimodal machine learning identifies low *ACO2* expression as a key predictor and determinant of the HP phenotype in clinical isolates. (A) Schematic of the multimodal data integration and machine-learning framework. (B) Bootstrap analysis of the representativeness of the 87-isolate subset. The histogram shows median AmB persistence values from 10,000 random resamplings of 87 clinical isolates from the 1,025-isolate collection; the dashed line marks the actual median persistence level of the selected subset. (C) ROC-AUCs for predicting HP clinical isolates using Random Forest and XGBoost models. (D) SHAP summary plots of the top 10 transcriptomic features for predicting HP clinical isolates using Random Forest and XGBoost (left), with corresponding mean absolute SHAP values indicating feature importance (right). (E) ROC curve for predicting HP clinical isolates using a single-variable Logistic Regression model based solely on *ACO2* expression. (F) *ACO2* expression levels in HP and NHP clinical isolates. (G) Percent survival of the SPCs of wild-type, *aco2*Δ, and *aco2*Δ + *ACO2* strains after 24-h treatment with AmB (10 × MIC). Data shown as mean ± SD (n = 3). (H) MIC evaluation of wild-type, *aco2*Δ, and *aco2*Δ + *ACO2* strains. (I) Percent survival of the SPCs of two HP clinical isolates with or without *ACO2* overexpression (OE) after 24-h AmB treatment. Data shown as mean ± SD (n = 3). (J) Proportions of *P_SPS1_-GFP::SPS1* and *aco2*Δ/*P_SPS1_-GFP::SPS1* strains in the cell populations with low Sps1 expression and high Sps1 expression after 7 days in mice with or without AmB treatment. Data shown as mean ± SD (n = 3). ns, not significant; \*\**p* < 0.01; \*\*\**p* < 0.001.

To identify the major transcriptomic driver of model prediction, we further assessed the contribution of each gene’s transcript level to the predictive models using both feature importance and SHAP (SHapley Additive exPlanations) values.^87, 88^ Under both interpretative frameworks, *CNAG_07908* emerged as the top-ranking feature (Figure 4D). This gene is annotated as the mitochondrial aconitase-encoding gene *ACO2*.^89, 90^ Notably, *ACO2* expression alone showed strong predictive power for HP variants, achieving an AUC of 0.955 (Figure 4E). SHAP analysis suggested a negative association between *ACO2* expression and the HP phenotype, and group-wise comparison confirmed that *ACO2* expression was significantly lower in HP clinical isolates than in NHP isolates (Figure 4F).

Given that low *ACO2* expression accurately predicts HP isolates, we hypothesized that *ACO2* expression deficiency may be a determinant underlying the HP phenotype in clinical *C. neoformans* isolates. To test this, we generated an *ACO2* deletion mutant (*aco2*Δ) in the reference strain H99 background. Loss of *ACO2* significantly increased the survival rate against AmB killing in SPCs by approximately one order of magnitude (Figure 4G). Furthermore, *ACO2* deletion did not alter MIC values (Figure 4H), indicating that *ACO2* is an AmB persistence-specific gene. Reintroduction of the native *ACO2* gene into the *aco2*Δ mutant reduced persistence to the wild-type levels (Figure 4G). These results demonstrate that *ACO2* significantly and negatively affects AmB persistence in *C. neoformans*. To further confirm the direct effect of *ACO2* expression on persistence levels in clinical isolates, we overexpressed *ACO2* in two HP clinical isolates from distinct genetic backgrounds. *ACO2* overexpression dramatically reduced the survival rate of SPCs upon AmB exposure by one to two orders of magnitude (Figure 4I). Together, these results demonstrate that the *ACO2* expression level is a key determinant of the HP phenotype in clinical isolates.

Moreover, we performed animal experiments to evaluate the effect of *ACO2* deficiency on AmB persistence *in vivo*. The *aco2*Δ/P*_SPS1_-GFP::SPS1* and P*_SPS1_-GFP::SPS1* strains were mixed at a 1:1 ratio and used for co-infection in mice as previously mentioned (Figure 2G). After 7 days of AmB treatment, the *aco2*Δ/P*_SPS1_-GFP::SPS1* strain was significantly enriched in the Sps1^high^ subpopulation compared to the P*_SPS1_-GFP::SPS1* strain, whereas no significant difference was observed in the Sps1^low^ subpopulation (Figure 4J). In untreated co-infected mice, no significant differences were detected between the two strains regardless of Sps1 expression level (Figure 4J). These results indicate that *ACO2* deficiency significantly enhances dormancy-mediated AmB persistence *in vivo* without incurring an apparent fitness cost.

### *ACO2* deficiency promotes POPC accumulation, which enhances AmB persistence by competing with AmB for ergosterol

Although annotated as a mitochondrial aconitase that may function in the tricarboxylic acid (TCA) cycle,^91, 92^ the detailed molecular function of Aco2 has not yet been validated in *C. neoformans*. We thus performed subcellular localization analysis using an Aco2-mCherry fusion expression system and confirmed its mitochondrial localization (Figure 5A).^93^ Furthermore, we heterologously expressed and purified *C. neoformans*-derived Aco2 protein in *Escherichia coli* (Figure S5A),^94^ and purified recombinant Aco2 displayed aconitase activity by catalyzing the conversion of isocitrate to citrate *in vitro* (Figure 5B).^95^ These results indicate that Aco2 functions as a mitochondrial aconitase in *C. neoformans*.

**Figure 5.**
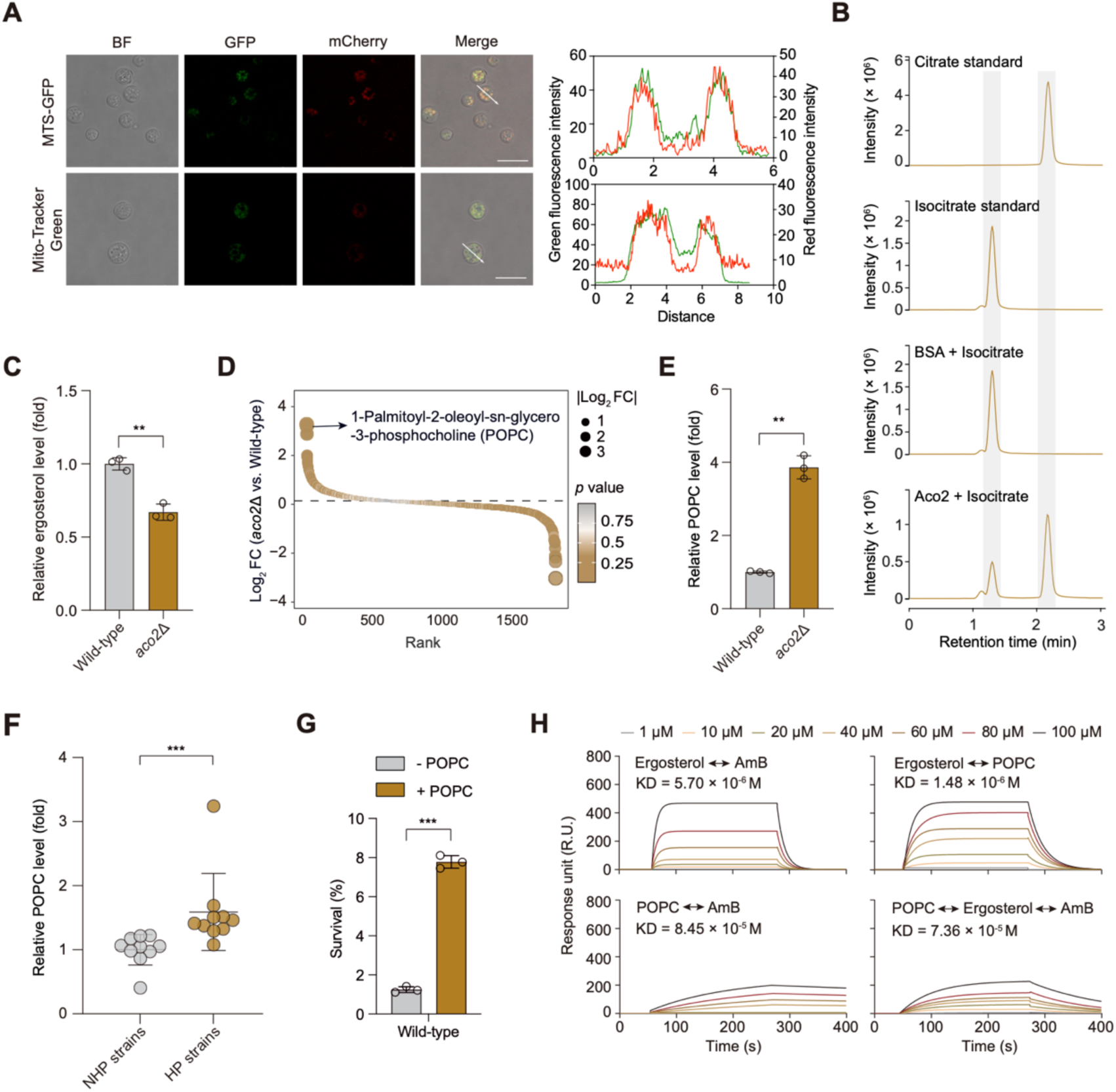
*ACO2* deficiency promotes POPC accumulation, which attenuates AmB killing by competing for ergosterol. (A) Colocalization of Aco2-mCherry with mitochondria marked by MTS-GFP or MitoTracker Green staining. Histograms of fluorescence intensity were plotted in the direction indicated by the white arrows. BF, bright field. Scale bars, 10 μm. (B) Extracted ion chromatogram (EIC) at *m/z* 191.0197 ([M-H]^-^) from LC-MS analysis of citrate and isocitrate standards and reaction mixtures containing isocitrate incubated with either BSA or Aco2. (C) Relative ergosterol levels of wild-type and *aco2*Δ SPCs. Data shown as mean ± SD (n = 3). (D) Rank of metabolites based on the metabolic profiling of wild-type and *aco2*Δ SPCs. (E) Relative POPC levels of wild-type and *aco2*Δ SPCs. Data shown as mean ± SD (n = 3). (F) Relative POPC levels of NHP and HP clinical isolates. Each point represents the mean value from three independent biological replicates. Data shown as mean ± SD. (G) Percent survival after 24-h treatment with AmB (10 × MIC) of H99 wild-type strain in the absence or presence of exogenous POPC. Data shown as mean ± SD (n = 3). (H) Sensorgrams for binding affinities between AmB, ergosterol and POPC, liposomes immobilized on the Lip-1 sensor chip surface. Data homogenization was done using GraphPad Prism 8. \*\**p* < 0.01; \*\*\**p* < 0.001.

Given that Aco2 is a mitochondrial aconitase involved in the TCA cycle, we hypothesized that its deficiency may lead to metabolic reprogramming that affects AmB persistence. To test this, we first used LC-MS to compare the abundance of the metabolite ergosterol, the target of AmB,^2, 36^ in SPCs of the *aco2*Δ mutant and the wild-type strain. The results showed that *ACO2* deletion significantly reduces ergosterol content, decreasing to 0.67-fold of WT levels (Figure 5C). However, this reduction appears insufficient to fully account for the nearly one-order-of-magnitude increase in survival upon AmB treatment in the *aco2*Δ mutant, suggesting the existence of parallel mechanisms.

To further investigate this, we conducted untargeted metabolomics to profile the metabolic landscape of SPCs from wild-type and *aco2*Δ mutants (Table S4). Among the differentially abundant metabolites, POPC (1-palmitoyl-2-oleoyl-sn-glycero-3-phosphocholine) showed the greatest upregulation in *aco2*Δ SPCs (Figure 5D). POPC production and physiological function in *C. neoformans* remain largely unclear. To validate the aforementioned metabolomics-based findings, we performed targeted quantification using a POPC standard combined with high-resolution LC-MS. This confirmed that POPC is present in *C. neoformans* SPCs (Figures S5B and S5C) and that its abundance is significantly higher in the *aco2*Δ mutant than in the wild-type strain (Figure 5E).^96, 97^ We further assessed the correlation between POPC content and persistence levels in clinical isolates and found that clinical HP strains had significantly higher POPC levels than NHP strains (Figure 5F).

The fact that both POPC and ergosterol are membrane components raises the hypothesis that POPC may interact with ergosterol on the membrane, which limits AmB binding to ergosterol and, as a result, limits AmB efficacy. To test this hypothesis, we added POPC exogenously to wild-type SPCs, and analyzed its effect on AmB persistence. Exogenous POPC supplementation significantly increased the survival of wild-type SPCs against AmB killing (Figure 5G). This indicates that increased POPC levels significantly promote AmB persistence in *C. neoformans*.

To verify the potential interaction between POPC and ergosterol that limits AmB efficacy, we performed Surface Plasmon Resonance (SPR) analysis.^36^ The results showed a strong interaction between POPC and ergosterol with a dissociation constant (*K*_D_) value of 1.48 × 10⁻⁶ M, with an affinity significantly higher than that between AmB and ergosterol with a *K*_D_ value of 5.70 × 10⁻⁶ M (Figure 5H). Importantly, in the presence of POPC, the binding affinity of AmB for ergosterol decreased by nearly an order of magnitude (Figure 5H), indicating that POPC can restrict AmB-ergosterol binding.

### The clinical-stage antifungal T-2307 effectively eliminates clinical cryptococcal HP strains irrespective of genetic background

The difficulty in clearing HP variants *in vivo* associated with *ACO2* expression deficiency prompted us to search for potential therapeutic agents effective against these refractory strains. We therefore screened a total of 1,935 compounds comprising a library of Traditional Chinese Medicine and clinical-stage anti-*Cryptococcus* candidates, including sertraline that has been demonstrated to effectively kill cryptococcal persisters,^57, 98^ for their ability to eliminate SPCs of the *aco2*Δ mutant (Figure 6A). Among the compounds tested, eight exhibited greater killing activity against *aco2*Δ SPCs compared with AmB (Figure 6B and Table S5). Notably, these compounds included chlorhexidine diacetate and tomatine, both previously reported to possess antifungal activity against *C. neoformans*.^99^ The presence of these known active agents validated the reliability of our screening platform.

**Figure 6.**
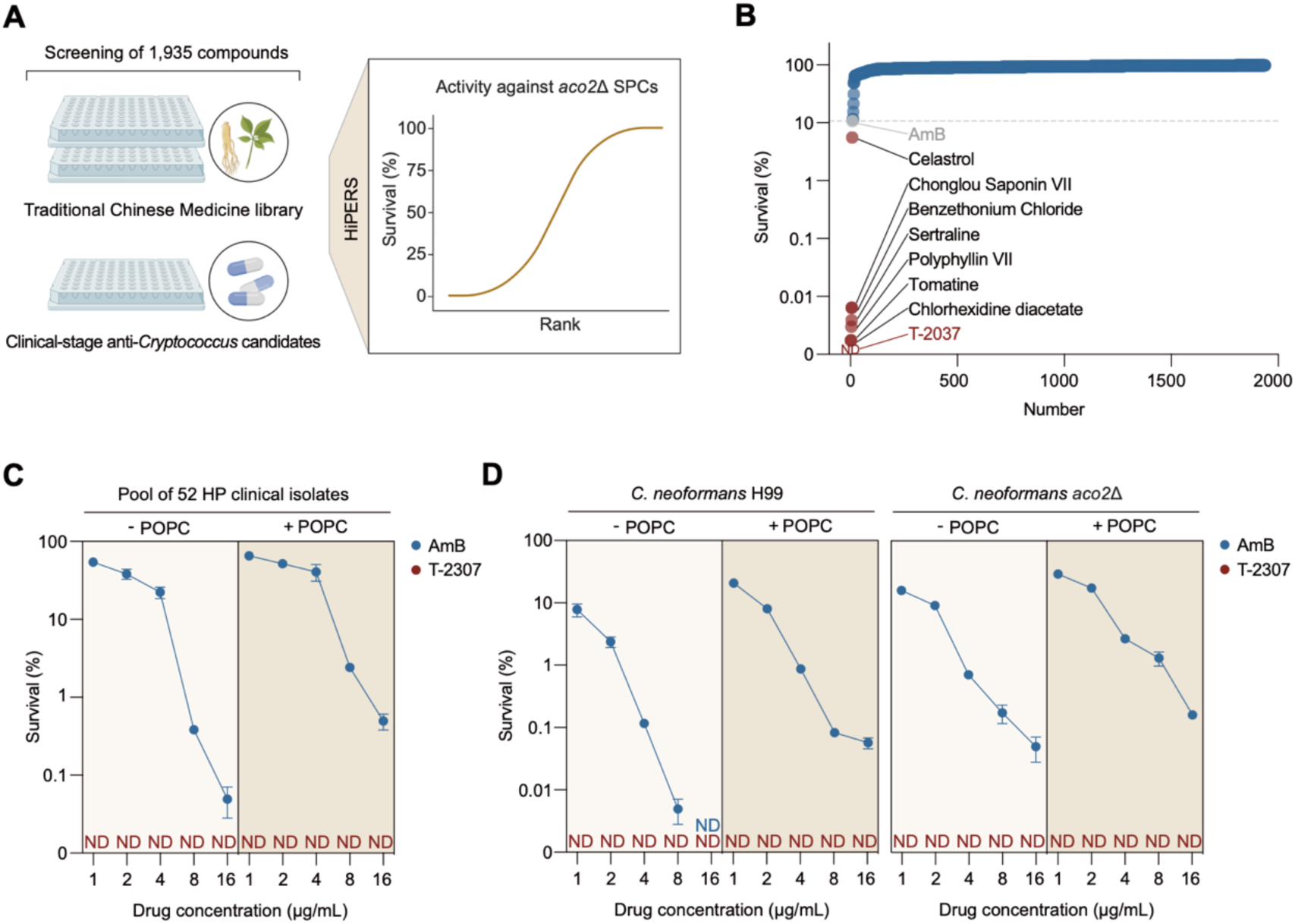
T-2307 efficiently eliminates high-persistence *C. neoformans* strains. (A) Schematic diagram of the experimental workflow to identify drugs acting against *aco2*Δ SPCs. (B) Ranked survival of *aco2*Δ SPCs across the 1,935-compound screen. Eight compounds exhibiting greater killing activity than AmB are highlighted in red. Each point represents the mean survival from two independent biological replicates. ND, not detected. (C) Percent survival of the SPCs of a pooled population of 52 HP clinical isolates after 24-h treatment with the indicated concentrations of AmB or T-2307, in the absence or presence of exogenous POPC. Data are shown as mean ± SD (n = 3). ND, not detected. (D) Percent survival of SPCs of wild-type and *aco2Δ* strains after 24-h treatment with the indicated concentrations of AmB or T-2307, in the absence or presence of exogenous POPC. Data are shown as mean ± SD (n = 3). ND, not detected.

Among the compounds tested, T-2307, an arylamidine antifungal candidate in the clinical development (as listed in the WHO 2025 antifungal preclinical and clinical pipeline review),^100–106^ exhibited the most potent activity against hyper-persistent *aco2*Δ mutant, outperforming even sertraline (Figure 6B). We further evaluated its killing efficacy against a pool of all 52 HP clinical isolates from the CHIF-NET collection, which represent diverse genetic backgrounds. Following treatment with 1 µg/mL T-2307 for 24 h, the viable SPCs from the pool of HP clinical isolates were reduced below the detection limit (Figure 6C). These results demonstrate that the potent activity of T-2307 is not restricted to the *aco2*Δ mutant constructed in the H99 reference background, but extends broadly to HP clinical isolates with diverse genetic lineages.

We next compared the fungicidal activities of T-2307 and AmB under conditions of exogenous POPC supplementation. Consistent with our previous findings, the addition of POPC markedly compromised the killing efficacy of AmB against SPCs of both wild-type and *aco2*Δ strains in the H99 background (Figure 6D). In contrast, exogenous POPC did not prevent T-2307-mediated killing under the conditions tested (Figures 6C and 6D). Together, these findings demonstrate that T-2307 shows promising efficacy against both *aco2*Δ SPCs and clinical HP strains, even in the presence of POPC, which promotes AmB persistence.

## DISCUSSION

In this study, we developed a high-throughput strategy for quantifying fungicide persistence and used it to profile a nationwide collection of clinical *C. neoformans* isolates from the CHIF-NET program. We observed marked variation in AmB persistence levels among clinical isolates (Figure 1C), which led to our finding that AmB persistence can readily evolve in patients with cryptococcosis (Figures 2C-2H). Functional genomics revealed that this persistence evolution is likely attributable to the minimal fitness costs associated with persistence-related mutations (Figures 3C and 3D). Future prospective cohort studies may help establish a direct correlation between persistence evolution and treatment outcomes in patients with cryptococcosis or other fungal diseases.^107^ In this regard, given the marked variation in persistence levels among isolates from the same patient, traditional examinations that test only a few isolates may introduce considerable bias, preventing the establishment of accurate correlations. Notably, analogous concerns in bacterial infection research have led to recommendations for population-level sampling of clinical specimens, rather than relying on one or a few single-colony isolates, to minimize trait-dependent biases in prognostic or resistance analysis.^50, 108, 109^ The HiPERS strategy, by enabling the assessment of hundreds of isolates per patient, allows evaluation of the overall persistence levels of *in vivo*-evolved populations and their association with treatment outcomes.

By integrating multimodal data with machine learning, we identified deficient expression of the mitochondrial aconitase gene *ACO2* as a key interpretable biomarker for HP clinical *C. neoformans* isolates (Figure 4E). Consistently, we demonstrated that *ACO2* deletion recapitulates the high-persistence phenotype by reprogramming lipid metabolism: it suppresses the synthesis of ergosterol (the target of AmB) while promoting the production of POPC, a phosphatidylcholine (PC) species that strongly competes with AmB for ergosterol binding (Figure 5H). Previous studies have shown that POPC is a conserved phospholipid component of cell membranes in eukaryotes, including mammals. ^110–114^ Moreover, several microbial pathogens have been shown to acquire PC from mammalian hosts.^115–117^ Given the strong effect of POPC on triggering cryptococcal persistence, this raises the possibility that mammalian host-derived POPC may also facilitate fungal survival against AmB treatment, a possibility that warrants future investigation.

Through drug screening, we identified T-2307 as the most potent agent against the hyper-persistent *aco2*Δ mutant among all compounds tested in this study (Figure 6B). This panel included sertraline, which has previously been reported to exhibit remarkable efficacy against cryptococcal persisters,^57^ thereby validating the reliability of the screening platform. T-2307 is a clinical-stage antifungal candidate under development for cryptococcosis, and previous studies in healthy volunteers have reported no drug-related serious adverse events.^106^ We further demonstrated that T-2307 displayed comparable killing efficacy against the pool of HP clinical isolates from the CHIF-NET collection, which encompasses diverse genetic lineages as revealed by whole-genome sequencing (Figure 6C). Collectively, these data support the potential of T-2307 for eliminating cryptococcal HP isolates in real-world clinical settings.

Overall, our study demonstrates that AmB persistence evolution can occur in cryptococcal patients and may represent a previously neglected clinical concern. Our findings also raise exciting questions: Does within-host evolution of fungal persistence represent a general mechanism underlying incomplete fungal clearance during fungicidal treatment across different invasive fungal infections? How do host–fungus– drug interactions drive the evolution of persistence traits during infection and treatment? Future work on these questions may foster therapies that enhance *in vivo* fungal clearance, thus improving cure rates across diverse fungal diseases.

## RESOURCE AVAILABILITY

### Lead contact

Further information and requests for resources and reagents can be directed to and will be fulfilled by the lead contact, Linqi Wang.

### Materials availability

There are no restrictions on the availability of materials. Strains and plasmids used or generated in this study are available from the lead contact upon request.

## ACKNOWLEDGMENTS

We are grateful to all participants in the CHIF-NET program. We are grateful to the laboratory of Dr. Hiten Madhani (University of California) for making their *C. neoformans* mutant libraries available to the community.

This study was financially supported by National Natural Science Foundation of China (32425007 [L.W.], 32461160269 [L.W.][K.H.W.], 82502745 [W.K.], 82472308 [X.F.]); National Key Research and Development Program of China (2025YFE0205600 [M.X.] [L.W.] [Y. Xu]); Strategic Priority Research Program of the Chinese Academy of Sciences (XDB0810000 [L.W.]); the Chinese Academy of Sciences Project for Young Scientists in Basic Research (YSBR-111 [L.W.]); and the Science and Technology Development Fund, Macao S.A.R (FDCT) (0142/2024/AFJ [K.H.W.]). The funders had no role in study design, data collection and analysis, decision to publish, or preparation of the manuscript.

## AUTHOR CONTRIBUTIONS

Conceptualization, Y. Xie and L.W.; Methodology, Y. Xie, W.K., X.F., H.X., W.W., Z.Z., N.Z., H.M., Z.T., H.Z., L.J., B.Z., G.S., S.Y., M.Y., F.L., X.W., S.L., J.X., X.X., C.L., M.X., and L.W.; validation, Y. Xie, W.K., X.F. and H.X.; formal analysis, Y. Xie, W.K., X.F., H.X., W.W., Z.Z., H.M., N.Z., Z.T., H.Z., L.J., B.Z., G.S., S.Y., M.Y., F.L. and X.W.; investigation, Y. Xie, W.K., X.F., H.X., W.W., Z.Z., H.M., N.Z., Z.T., H.Z., L.J., B.Z., G.S., S.Y., M.Y., F.L. and X.W.; funding acquisition, W.K., K.H.W., X.F., Y. Xu, M.X., and L.W.; resources, X.F., H.X., W.W., B.Z., F.L., X.W., J.X., C.H.L., X.X., Y. Xu, C.L., M.X., and L.W.; writing for original draft, Y. Xie, W.K. and L.W.; writing for review & editing, Y. Xie, W.K, X.F., H.X., K.H.W., C.H.L., D.S.P., Y. Xu, C.L., M.X., and L.W.; visualization, Y. Xie, W.K., Z.Z., N.Z., H.M., Z.T., H.Z., L.J., Y. Xu, C.L., M.X. and L.W.

## DECLARATION OF INTERESTS

The authors declare no competing interests.

## EXPERIMENTAL MODEL AND STUDY PARTICIPANT DETAILS

### Strains and growth conditions

All strains used in this study are listed in Table S1 and S6. A total of 1,025 clinical *C. neoformans* isolates were collected through the CHIF-NET program and the research use of these clinical isolates was approved by the Peking Union Medical College Hospital Ethics Committee (JS-3009).

*C. neoformans* strains were cultured in yeast extract peptone dextrose (YPD) liquid medium (2% glucose, 2% peptone, and 1% yeast extract) at 30°C with shaking at 220 rpm, unless otherwise indicated. *E. coli* strains were cultured in Luria-Bertani (LB) broth or on LB agar plates supplemented with specific antibiotics at 37°C. MIC testing was performed using RPMI 1640 medium buffered with morpholinepropanesulfonic acid (MOPS), according to CLSI guidelines. E-test assays were performed on solid RPMI 1640 medium (MOPS-buffered and supplemented with 2% glucose and 1.5% agar), which were incubated at 35°C in the dark for 72 h, following established protocols.^118^

### Mice

All mouse experiments were performed under the guidance of “the regulation of the Institute of Microbiology, Chinese Academy of Sciences of Research Ethics Committee”. The mouse models and procedures performed have been approved by the Institute of Microbiology, Chinese Academy of Sciences of Research Ethics Committee (Permit No. SQ-APIMCAS2025036). Female C57BL/6 mice, 7-8 weeks old, were obtained from Vital River (Beijing, China). Mice were maintained under specific pathogen-free conditions in ventilated cages in a temperature- and humidity-controlled facility at 21°C and 50%-70% relative humidity, with a 12 h light/12 h dark cycle and unlimited food and water.

### Human subjects

Experiments with human CSF samples were approved by the ethics committees of the First Affiliated Hospital of Xiamen University (2026-163) and the First Affiliated Hospital of Nanchang University ((2025)CDYFYYLK(04-020)). Informed consent was waived for these excess samples. The CSF samples from five patients with cryptococcosis were included: three male patients aged 32, 53, and 65 years and two female patients aged 67 and 72 years.

## METHOD DETAILS

### Strain construction

To construct gene deletion strains in *Cryptococcus*, the nourseothricin (NAT) selectable marker cassette was fused to approximately 1.2-1.5 kb 5′ and 3′ homologous flanking regions amplified from the respective wild-type genomic templates. The resulting linear amplicons were introduced into designated *Cryptococcus* recipient strains via the transient CRISPR-Cas9 expression (TRACE) method, as previously described.^119^

For complementation, target open reading frames (ORFs) with their native promoter regions were PCR-amplified and subcloned into the pXC plasmid vector. For overexpression, target ORFs were amplified, subjected to appropriate restriction enzyme digestion, and cloned downstream of the constitutive *H3* promoter within the pXC vector. The constructs were subsequently integrated into the second safe haven (SH2) locus of the corresponding recipient backgrounds via electroporation.^120, 121^ GFP-Sps1 reporter strains were constructed as previously described.^57^ All primers utilized for strain construction in this study are listed in Table S7.

### MIC assay

The microdilution assay was performed according to the standard protocol of CLSI.^118^ *Cryptococcus* cells were diluted to 2.5 × 10^3^ cells/mL, and then AmB was added to the indicated concentrations. The MIC value was determined by comparing the growth of AmB-treated samples after 72 h of incubation with that of untreated controls by measuring the absorbance at 600 nm. For the E-test assay, cells were plated on solid RPMI 1640 medium. When the surface of the plate was completely dry, AmB-containing E-test strips were placed on each plate. The MIC value was read after 72 h of incubation at 35 °C according to the manufacturer’s instructions.^118^

### HiPERS methodology

To evaluate the killing efficacy of AmB, *C. neoformans* cells were inoculated into YPD liquid medium at an initial optical density (OD) of 1.0 in 96-well deep-well plates and cultured at 30°C with shaking (220 rpm) for 72 h to generate SPCs. One sterile 4.5-mm glass bead was added to each well to enhance agitation and maintain cells in suspension during cultivation. The resulting SPCs were harvested, resuspended in their original culture medium, and subsequently challenged with AmB (exceeding the CLSI ECV by more than 10-fold) at 37°C for 24 h, unless otherwise indicated. To assess the impact of POPC on the fungicidal activity of AmB, *C. neoformans* SPCs were supplemented with exogenous POPC lipid vesicles prepared as previously described and subsequently challenged with AmB at 37°C for 24 h.^122, 123^

Following AmB treatment, cell suspensions were serially diluted, and 5 μL of each dilution was spotted onto drug-free YPD agar plates using an automated platform. Percent survival was determined by calculating the ratio of colony-forming units (CFUs) between AmB-treated samples and untreated controls, with colonies enumerated after 2 days of incubation at 30°C.

To enable high-throughput colony quantification, an automated image-analysis pipeline was developed. Raw images of square YPD agar plates containing 96-position colony arrays were initially processed using Hough circle transform and perspective transformation algorithms to correct geometric distortions. Each array position was then cropped into an individual sub-image for batch processing. Sub-images were enhanced by using CLAHE and a sharpening filter to improve colony-background contrast and colony-edge definition. Colonies were subsequently segmented using SAM 2.^67^ Each detected colony was assigned a unique segmentation mask, and the number of masks per well was automatically recorded for CFU quantification.

### Genome sequencing and assembly

Genomic DNA was isolated utilizing a QIAamp DNA Mini Kit (Qiagen, Hilden, Germany) according to the manufacturer’s instructions.^71^ A paired-end library was prepared and sequenced on the Illumina HiSeq PE150 platform. Raw short-read FASTQ files were quality-filtered through a standardized Trimmomatic (v0.39) pipeline and FastUniq (v1.1) was used to remove PCR duplicates.^124^ The quality of the filtered reads was assessed using FastQC (v0.11.5). De novo genome assembly was performed using SPAdes (v3.13.0).^125^ The quality of the genome assembly was assessed using QUAST (v5.3.0).^126^

### Variant calling

The quality-filtered reads were aligned to the *C. neoformans* H99 reference genome (GenBank assembly accession: GCA_000149245.3) using bwa mem,^127^ and BAM files were generated with SAMtools (v1.6).^128^ Variants were called with GATK (v4.3.0.0) using HaplotypeCaller.^129^ After calling, variants were filtered on the following parameters: QD < 2.0, QUAL < 30.0, SOR > 3.0, FS > 60.0 (InDels > 200), MQ < 40.

### MLST analysis

MLST analysis was performed according to the ISHAM consensus scheme using the seven loci *CAP59, GPD1, LAC1, PLB1, SOD1, URA5,* and *IGS1*.^130^ The corresponding locus sequences were extracted from the genome assemblies and then queried in the official *Cryptococcus* MLST database (https://mlst.mycologylab.org/) for allele assignment and sequence type (ST) determination.^130^

### Phylogenetic analysis

To elucidate the phylogenetic relationships among the 1,025 *C. neoformans* clinical isolates, high-confidence SNPs from the multi-sample VCF were converted into FASTA format using vcf2phylip.py.^131^ A Maximum Likelihood (ML) phylogenetic tree was inferred using IQ-TREE (v3.0.1) with the optimal nucleotide substitution model estimated by ModelFinder Plus.^132^ Branch supports were assessed using 1,000 ultrafast bootstrap replicates.^133^

### Population structure analysis

Population structure was assessed using ADMIXTURE (v1.3).^134^ The number of components K was optimized to minimize the cross-validation error. PCA was independently performed on the same variant set using PLINK (v1.9).^135^

### Comparative fitness landscape analysis of AmB persistence- and resistance-associated deletion mutants in *C. neoformans*

To compare the fitness profiles of AmB persistence- and resistance-associated mutants, we leveraged the fitness dataset of the genome-wide deletion mutants reported by Boucher *et al*.^79^ In that study, the *C. neoformans* deletion mutant library was systematically profiled across 141 *in vitro* growth conditions and one *in vivo* condition, generating a comprehensive fitness landscape. After cross-border transportation, we successfully revived over 2,600 mutants from this library. Mutants were ranked by the previously reported fitness scores under AmB treatment, and the top 10% were defined as the AmB-resistant group.^79^ To specifically identify persistence-associated mutants, we excluded mutants with significant effects on AmB resistance using the previously established threshold (|fitness score| ≥ 2).^79^ Among the remaining mutants, the top 10% ranked by survival following AmB exposure in HiPERS were defined as the AmB-persistent group. Fitness profiles of AmB persistence- and resistance-associated mutants were subsequently compared across 141 non-AmB conditions.

### Quantification of ergosterol by LC-MS

Ergosterol quantification was performed as previously described,^36, 136^ with slight modifications. In brief, *Cryptococcus* SPCs were washed twice with ice-cold (4°C) sterile water. Following lyophilization, cells were disrupted with ceramic beads and then suspended in chloroform supplemented with 2.5 μg/mL ketoconazole as an internal standard. The homogenate was agitated for 8 h, after which the supernatant was collected and dried. Methanol:chloroform (10:1, v/v) was used to redissolve the sample for later LC–MS analysis. To prevent light-mediated degradation, all extraction procedures were strictly conducted in the dark.

Chromatographic separation was performed on an Agilent 1260 Series LC system equipped with an Agilent Eclipse Plus C18 column (1.8 μm, 2.1 × 50 mm) maintained at 45°C.^137^ The mobile phase consisted of water containing 0.1% acetic acid (solvent A) and acetonitrile (solvent B), delivered at a flow rate of 0.4 mL/min. The elution program was conducted as follows: 0-2 min, 70%-100% B; 2-18 min, 100% B; 18-19 min, 100%-70% B; and 19-24 min, 70% B for column re-equilibration.

Mass spectrometric analysis was conducted using an Agilent Accurate-Mass Q-TOF 6520B instrument in positive ionization mode with an atmospheric pressure chemical ionization (APCI) source. Key instrumental parameters were set as follows: fragmentor voltage, 130 V; capillary voltage, 3,500 V; corona current, 4 μA; nebulizer pressure, 50 psi; drying gas and vaporizer temperature, 350°C; and nitrogen gas (nebulizing and drying) flow rate, 4 L/min. Full-scan mass spectra were acquired over a mass range of mass-to-charge (*m/z*) 80-1,000 at an acquisition rate of 1.03 spectra/s. The final ergosterol concentrations were determined by normalizing the calculated amounts to the initial biomass of each sample.

### Quantification of POPC by LC-MS

POPC extraction was performed as previously described,^138^ with minor modifications. In brief, lyophilized and disrupted *Cryptococcus* SPCs were suspended with methanol containing 5 μg/mL 1,2-dimyristoyl-sn-glycero-3-phosphocholine (DMPC) as an internal standard. Methyl tert-butyl ether (MTBE) was added, and the samples were agitated for 2 h. Following the addition of ultrapure water, the samples were centrifuged to induce phase separation, and the upper organic phase was collected and dried. Methanol:isopropanol:water (65:30:5, v/v/v) was used to redissolve the resulting lipid extracts for LC-MS analysis.

Chromatographic separation was performed on an ultra-high-performance liquid chromatography (UHPLC) system equipped with a Waters ACQUITY^TM^ UPLC BEH C8 column (1.7 μm, 2.1 × 100 mm) maintained at 55°C. The samples were held at 4°C. The mobile phase consisted of acetonitrile:water (60:40, v/v; solvent A) and isopropanol:acetonitrile (90:10, v/v; solvent B), both supplemented with 10 mM ammonium acetate, and delivered at a flow rate of 0.26 mL/min. The gradient elution program was configured as follows: 0-1.5 min, 32% B; 1.5-15.5 min, 32%-85% B; 15.5-15.6 min, 85%-97% B; 15.6-18.0 min, 97% B; 18.0-18.1 min, 97%-32% B; and 18.1-20.0 min, 32% B.

Mass spectrometric analysis was conducted using a Waters Xevo^TM^ MRT instrument in positive ionization mode with an electrospray ionization (ESI) source, with full-scan data acquired over a mass range of *m/z* 50-2,000 at an acquisition rate of 2 spectra/s. The source parameters were as follows: capillary voltage, 2,500V; cone voltage, 30V; desolvation temperature, 500°C; source temperature, 120°C; desolvation gas flow, 750 L/h and cone gas flow, 20 L/h.^139^ The final POPC concentrations were determined by normalizing the calculated amounts to the initial biomass of each sample.

### SPR analysis

Molecular interactions among POPC, ergosterol, and AmB were systematically evaluated via SPR spectroscopy using an OpenSPR system (Nicoya Lifesciences).^36, 140^ Briefly, liposomes containing POPC, ergosterol, or POPC-saturated ergosterol were prepared by thin-film hydration method and immobilized onto Lip-1 sensor chips (Nicoya Lifesciences). Serial dilutions of AmB or POPC were injected over the sensor surface at a flow rate of 20 μL/min at 25°C, with PBS (pH 7.4) containing 2.5% DMSO or 2.5% methanol used as the respective running buffers. Blank liposome served as the negative control. Binding assays were performed in two biological replicates, and the sensor surface was regenerated with 20 mM 3-((3-cholamidopropyl) dimethylammonio)-1-propanesulfonate (CHAPS) after each test.

### Matrix-assisted laser desorption ionization-time of flight mass spectrometry (MALDI-TOF MS) acquisition

MALDI-TOF MS assay was performed as previously described,^141^ with minor modifications. Harvested *Cryptococcus* SPCs were washed twice, resuspended in ultrapure water, and mechanically disrupted with ceramic beads. Absolute ethanol was added to the resulting homogenates, followed by centrifugation, and the resulting pellets were air-dried. The pellets were extracted with 70% formic acid followed by acetonitrile, and the resulting mixtures were centrifuged to collect the protein-containing supernatants for MALDI-TOF MS analysis.

For MALDI-TOF MS acquisition, 1 μL of each protein extract was spotted onto a clean MALDI target plate and air-dried. Each sample spot was then overlaid with 1 μL of matrix solution comprising saturated α-cyano-4-hydroxycinnamic acid (CHCA) dissolved in 50% acetonitrile and 2.5% trifluoroacetic acid (TFA), and dried completely to facilitate co-crystallization. Mass spectra were acquired using an EXS 3000 MALDI-TOF MS System (Zybio) across a *m/z* range of 2,000–20,000, with data acquisition performed using EX-Accuspec software (v3.3.4.8). *E. coli* ATCC 25922 was included in each analytical run as a quality-control reference, and the assay was considered valid only when this reference strain yielded a high-confidence identification score.

### MALDI-TOF MS data preprocessing

Data preprocessing of mass spectra was performed as previously described,^142^ with minor modifications. Briefly, mass spectra were preprocessed using the R package MALDIquant (version 1.22.3).^143^ Subsequently, spectra were subjected to square-root transformation, Savitzky-Golay smoothing, baseline correction using the SNIP algorithm (20 iterations), and total ion current (TIC) normalization, followed by restriction to an *m/z* range of 2,000-20,000. The resulting spectra were partitioned into 6,000 non-overlapping 3-Da bins, and intensities within each bin were summed to generate a fixed-length feature vector for each spectrum. The binned feature vectors from two replicates were averaged to generate a single sample-level feature representation for downstream machine learning analysis.

### High-throughput single-cell Raman spectra (SCRS) acquisition

Single-cell Raman flow cytometry was performed as previously described, ^144^ with minor modifications. In brief, to minimize invalid measurement, cultured cells from each designated strain were harvested, washed extensively, and resuspended to a final density of 1.0 × 10⁷ cells/mL. The prepared suspension was continuously infused into a microfluidic chip integrated with a positive dielectrophoresis-induced deterministic lateral displacement (pDEP-DLD) module. To achieve precise single cell alignment, a modulated pDEP-DLD force, driven by a function generator, was applied to focus and trap the cells precisely at the laser point.

SCRS acquisition was performed on a FlowRACS instrument (Qingdao Single-cell Biotech, CN) equipped with a Nd:YAG 532 nm laser emitter. The incident laser beam was precisely focused onto the trapped cells through a 50 × objective lens (NA = 0.7), with the laser power maintained at 200 mW. For high-throughput spectral acquisition, the integration time was set to 500 ms per cell. Under these parameters, full-spectrum SCRS data were continuously collected from 200 individual cells for each sample.

### SCRS data preprocessing

SCRS were preprocessed using the RamEx R package (version 1.0.0) as previously described, including baseline correction, Savitzky-Golay smoothing, and peak area normalization.^145^ Spectra from 200 single cells per sample were subsequently averaged to obtain a mean Raman spectrum, which was used as the sample-level input for downstream machine-learning analysis.

### Machine learning framework

To classify clinical HP strains, Random Forest and XGBoost classifiers were implemented using scikit-learn and xgboost packages, respectively.^146–148^ Input features were derived from preprocessed multimodal data: genomics, transcriptomics, mass spectra, and Raman spectra. For the genomic data preprocessing, the filtered dataset was first imputed using Beagle (v5.5) to infer missing genotypes,^149^ then converted into binary numerical encoding using PLINK and used as input features.^150^ For multimodal data integration, a feature-level early fusion strategy was executed by horizontally concatenating preprocessed feature sets from pairwise modalities into a unified input matrix. Total ten configurations were evaluated, including four unimodal and six pairwise feature sets.^151^ Both Random Forest and XGBoost classifiers were systematically trained across all configurations.

Model performance was assessed using five repeats of nested stratified fivefold cross-validation.^152^ Within the inner cross-validation loop, hyperparameter optimization was performed via Bayesian optimization using BayesSearchCV from the scikit-optimize package (version 0.10.2) over 25 iterations.^153^ The best-performing models were selected based on AUC-ROC scores from the inner cross-validation loop.

Model interpretability and feature attribution were established through SHAP analysis to quantify the relative contribution of individual predictors and identify key predictors of classification. Based on the SHAP analysis, a logistic regression model was constructed using *ACO2* expression as the predictor to evaluate its performance in predicting the clinical HP strains.

### RNA purification, RNA-seq and data analysis

Total RNA extraction of *Cryptococcus* SPCs was performed using the Ultrapure RNA Kit (Kangweishiji, CW0581S) as previously described.^154^ RNA levels and integrity were assessed by Qubit RNA Assay Kit in Qubit 2.0 Fluorometer (Life Technologies, CA, USA) and RNA Nano 6000 Assay Kit of the Bioanalyzer 2100 system (Agilent Technologies, CA, USA), respectively. RNA purity was evaluated using the Nano Photometer spectrophotometer (IMPLEN, CA, USA).

Transcriptome libraries were constructed utilizing the VAHTS mRNA-seq v2 Library Prep Kit (Vazyme Biotech). Subsequently, samples were clustered using VAHTS RNA adapters (Set 1/Set 2) and sequenced in a 2 × 150 pair-ended manner. RNA sequencing was performed on an Illumina platform by Co., Ltd (Beijing, China). Initial quality control of the sequencing data was performed using FastQC (v0.11.5). Approximately 2 GB of clean data per sample were mapped to the annotated *C. neoformans* H99 reference genome using HISAT2 (v2.2.1). Gene expression levels were quantified as transcripts per million (TPM) utilizing featureCounts (v2.1.1).

### Metabolomics

SPCs from the wild-type and *aco2*Δ strains of *C. neoformans* were harvested, and suspended in ice-cold methanol:water (4:1, v/v) containing an internal standard. Following three freeze-thaw cycles, extracts were centrifuged and the supernatants were collected for LC-MS analysis. Equal volumes of metabolites from each sample were mixed to generate quality control samples, which were analyzed at regular intervals to assess the reproducibility of the analytical procedure.

Metabolites were separated on a Waters ACQUITY^TM^ Premier HSS T3 column (1.8 μm, 2.1 × 100 mm) maintained at 45°C, with a flow rate of 0.4 mL/min. Water and acetonitrile, both containing 0.1% formic acid, were applied as mobile phases A and B respectively. MS was performed using electrospray ionization in the positive and negative ion scanning mode over *m/z* 70-1,000 at 60,000 resolution. Metabolomics data analysis was performed as previously reported.^57^

### *In vivo* model of systemic fungal infection

Mouse infections were performed as previously described,^57, 75^ with minor modifications. In brief, 7 to 8-week-old female C57BL/6J mice (Vital River, Beijing, China) were anesthetized with 30% (v/v) isoflurane diluted in propylene glycol prior to cryptococcal infection.

To conduct *in vivo* experimental evolution, mice were intranasally inoculated with 50 μL of a fungal suspension (2.0 × 10^6^ cells/mL) consisting of either the reference strain *C. neoformans* H99, a FLC^R^ clinical isolate, a 5-FC^R^ clinical isolate, or the hypervirulent *znf2*Δ mutant (H99 background). At 7 dpi, lung tissues were harvested and homogenized. The resulting homogenates were then plated onto YPD agar to recover individual fungal clones.

To test *in vivo* fungal burdens, mice (n = 3 per group) were randomly assigned and intranasally inoculated with 50 μL of a fungal suspension (2.0 × 10^6^ cells/mL). At 14 dpi, the lungs were harvested and homogenized in PBS. The resulting homogenates were serially diluted and plated onto YPD agar supplemented with 100 μg/mL chloramphenicol. Fungal burdens were quantified by enumerating CFUs following a 48-h incubation at 30°C.

For murine co-infection experiments with clinical strains, three independent pairs of matched within-patient high- and low-persistence isolates, each engineered with either a G418^R^ or Hyg^R^ marked P*_SPS1_*-*GFP*::*SPS1* reporter system, were mixed at a 1:1 ratio. For genetic validation, the *aco2*Δ/P*_SPS1_-GFP::SPS1* and P*_SPS1_-GFP::SPS1* strains were similarly mixed at a 1:1 ratio. In both experimental settings, mice were intranasally inoculated with 50 μL of the respective mixed fungal suspension, prepared from EEPCs, at a density of 2.0 × 10^8^ cells/mL. Following infection, mice (n = 3 per group) were randomly assigned to either a vehicle control or an AmB treatment group. AmB was administered intravenously at a dose of 1.0 mg/kg/day for 7 consecutive days, whereas the control group received an equivalent volume of 5% glucose vehicle via the same route.

### FACS isolation of fungal cells

FACS isolation of fungal cells was performed as previously described,^57^ with minor modifications. In brief, mice were intranasally inoculated with EEPCs of the designated reporter strains. To isolate Sps1^low^ and Sps1^high^ cell sub-populations, the lungs of infected mice were harvested and homogenized in 0.9% NaCl. Tissue homogenates were passed through a 40-μm cell strainer to prevent fluidic clogging and centrifuged at 4,000 rpm for 8 min. Following supernatant removal, pellets were resuspended in 0.9% NaCl, and *Cryptococcus* cells were stained with the cell wall dye Fluorescent Brightener 28 (Calcofluor White, 10 μg/mL) at 30°C for 10 min. Samples were then subjected to flow cytometry (FCM), with *Cryptococcus* cells explicitly gated based on Calcofluor White fluorescence. Cells were sorted on a BD FACSAria III flow cytometer to collect the GFP-Sps1^low^ (lowest 5% of the total population) and GFP-Sps1^high^ (highest 5% of the total population) fractions. Data analysis was performed using FlowJo software (TreeStar).

### Subcellular localization analysis of Aco2

Subcellular localization analysis was performed as previously described,^93^ with minor modifications. To determine the intracellular distribution of Aco2, an *ACO2*-mCherry fusion strain was constructed following the strain construction procedures described above. Mitochondrial localization of Aco2-mCherry was assessed by co-localization with either the mitochondrial targeting sequence GFP reporter (MTS-GFP) or MitoTracker Green.

Images were acquired on a Zeiss Axioplan 2 imaging system equipped with an AxioCam MRm camera using ZEN 2011 software (Carl Zeiss). High-resolution imaging was performed using a Leica TCS SP8 STED microscope, and images were processed with ImageJ (v1.53k).

### Aco2 expression and purification from *E. coli*

The intron-free coding sequence (CDS) of *C. neoformans ACO2* was cloned into the pET-22b(+) vector to generate the expression construct pET22b-*ACO2*-6xHis, which was subsequently transformed into *E. coli* BL21(DE3) competent cells for heterologous expression. *E. coli* BL21(DE3) were grown in LB broth supplemented with 100 µg/mL ampicillin until OD_600_ reached approximately 1.0. Protein expression was then induced by the addition of 0.5 mM isopropyl β-D-1-thiogalactopyranoside (IPTG) at 25 °C overnight.

Pellet was collected by centrifugation and resuspended in PBS supplemented with 0.1 mM phenylmethanesulfonyl fluoride (PMSF) and 0.1 mg/mL DNase I, followed by cell disruption via ultrasonication on ice. The lysate was clarified by centrifugation, and the resulting supernatant was loaded onto a 5-mL HisTrap Ni-NTA column (GE Healthcare). The His-tagged Aco2 protein was eluted using an imidazole gradient in a basal buffer (50 mM Tris-HCl, pH 7.8, 150 mM NaCl) according to the manufacturer’s protocol. The eluted protein was further purified by using the AKTA Explorer FPLC system with a Superdex 200 Increase 10/300 GL column. Protein purity and molecular identity were verified via SDS-PAGE, while the final protein concentration was quantified utilizing the Bradford protein assay.

### Iron-sulfur cluster reconstitution and aconitase activity measurement

The [4Fe–4S] cluster reconstitution of purified Aco2 and aconitase activity measurement were performed as previously described,^95^ with minor modifications. Briefly, the purified protein was incubated for 2 h in a reconstitution buffer (25 mM Tris-HCl, pH 7.4, 100 mM NaCl) supplemented with 75 μM (NH_4_)_2_Fe(SO_4_)_2_, 75 µM sodium hydrosulfide (NaHS) and 10 mM dithiothreitol (DTT). Following reconstitution, aconitase activity was assessed using isocitrate as the substrate. Reconstituted Aco2 (40 nM) was incubated with isocitrate (0.5 mM) overnight, whereas bovine serum albumin (BSA, 40 nM) was substituted for Aco2 in parallel negative-control reactions. The reaction mixture was analyzed by high-resolution LC-MS to detect the formation of citrate, thereby evaluating the catalytic activity of Aco2. All reconstitution procedures and enzymatic assays were rigorously performed under the aforementioned argon-saturated atmosphere to minimize oxidative degradation of the iron-sulfur cluster.

Chromatographic separation was performed on an UHPLC system equipped with a Waters ACQUITY^TM^ Premier HSS T3 column (1.8 μm, 2.1 × 100 mm) maintained at 40 °C, with a flow rate of 0.3 mL/min. Water containing 0.1% formic acid and acetonitrile were applied as mobile phases A and B respectively. The gradient elution program was set as follows: 0-1 min, 100% A; 1-5 min, 100%-90% A; 5-6 min, 90%-0% A; 6-11 min, 0% A; 11-16 min, 0%-100%A.^155^

MS analysis was conducted using a Waters Xevo^TM^ MRT instrument in negative ionization mode with an ESI source, with full-scan data acquired over a mass range of *m/z* 50-1,200 at an acquisition rate of 1 spectrum/s. The source parameters were set as follows: capillary voltage, 2,000 V; cone voltage, 40 V; desolvation temperature, 450 °C; source temperature, 120 °C; desolvation gas flow, 800 L/h and cone gas flow, 50 L/h.

## QUANTIFICATION AND STATISTICAL ANALYSIS

All quantitative data are presented as the mean ± SD, unless otherwise specified in the corresponding figure legends. Statistical parameters, including the exact value and definition of n, the statistical tests used, and significance thresholds, are provided in the figure legends or Method Details.

Statistical analysis were primarily executed using the R statistical computing environment (v4.5.1) and GraphPad Prism 8, unless otherwise specified in figure legends or Method Details. Correlations between quantitative variables were assessed using Pearson’s correlation coefficient, with Pearson’s *r* and corresponding *p* values reported where applicable. Pairwise comparisons between two independent groups were analyzed using a two-tailed, unpaired Student’s t-test. Comparisons of AmB persistence levels across genetic clusters were analyzed using Kruskal-Wallis tests. For image-analysis benchmarking, model performance and counting accuracy were evaluated using R^2^, Pearson’s *r*, MAE, and RMSE. For machine-learning analysis, classification performance was evaluated by ROC-AUC, and feature contributions were interpreted using feature-importance scores and SHAP values as described in Method Details. A *p* value of < 0.05 was considered statistically significant unless otherwise indicated. Statistical significance is denoted in the figures as follows: * *p* < 0.05, ** *p* < 0.01, *** *p* < 0.001, and ns (not significant).

---

**Figure S1.**
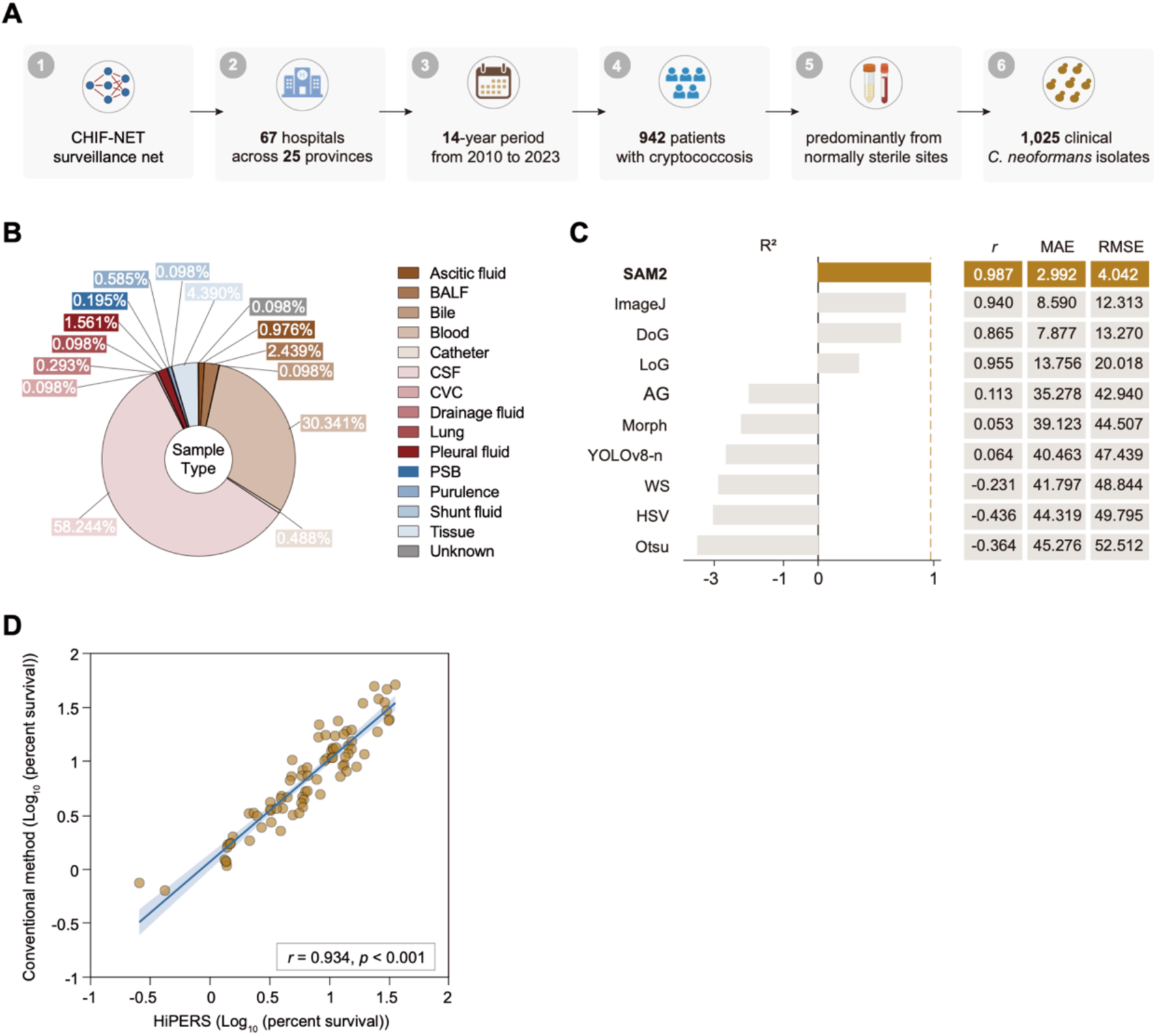
Nationwide collection of clinical *C. neoformans* isolates and validation of HiPERS, related to Figure 1. (A) Schematic overview of sample collection through the CHIF-NET program. The figure was generated in BioRender. (B) The proportions of clinical *C. neoformans* isolates from sites of isolation. BALF, Bronchoalveolar lavage fluid; CSF, Cerebrospinal fluid; CVC, Central Venous Catheter; PSB, Protected Specimen Brush. (C) Comparison of SAM 2-based colony counting with conventional image-analysis approaches. The bar plot shows the coefficient of determination R² relative to manual counting; the accompanying table summarizes Pearson’s *r*, MAE, and root mean square error (RMSE). DoG, Difference of Gaussian; LoG, Laplacian of Gaussian; AG, adaptive Gaussian thresholding; Morph, morphology-based segmentation; YOLOv8-n, You Only Look Once version 8 nano; WS, watershed segmentation; HSV, hue-saturation-value color-space thresholding; Otsu, Otsu’s automatic thresholding method. (D) Correlation between AmB persistence measurements obtained by HiPERS and the conventional method. Pearson’s *r* and *p* value are indicated.

**Figure S2.**
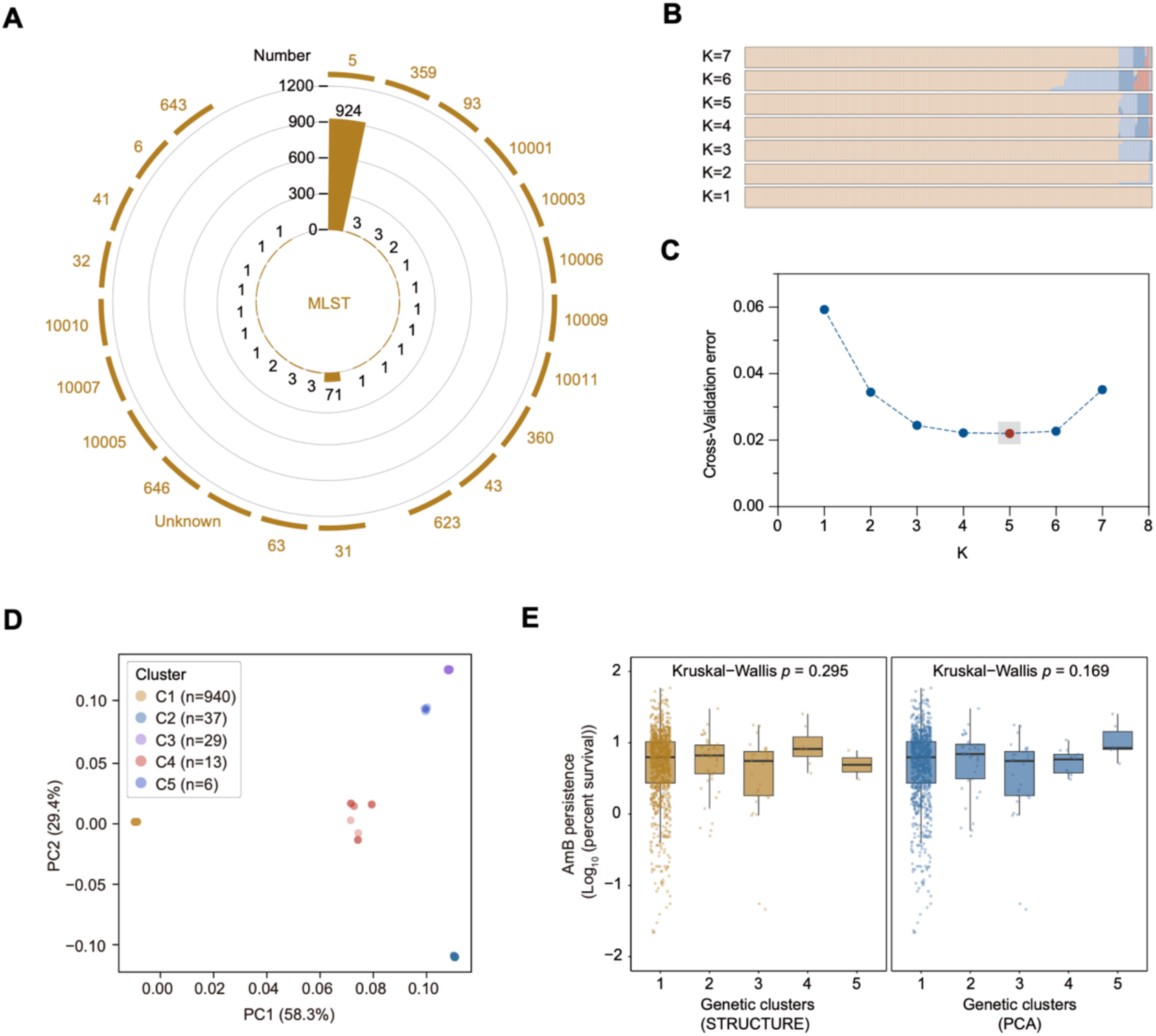
AmB persistence varies within genetic clusters of clinical *C. neoformans* isolates but does not differ significantly across clusters, related to Figure 1. (A) Distribution of MLST sequence types among clinical *C. neoformans* isolates. (B) STRUCTURE analysis of clinical *C. neoformans* isolates for K = 1 to 7.. (C) Cross-validation error for different values of K, with a minimum at K = 5. (D) PCA based on whole-genome SNPs of clinical *C. neoformans* isolates. The positions of the cluster of clinical isolates are indicated, each distinguished by a unique color. (E) AmB persistence levels of clinical *C. neoformans* isolates across genetic clusters defined by STRUCTURE or PCA. Kruskal-Wallis *p* values are indicated.

**Figure S3.**
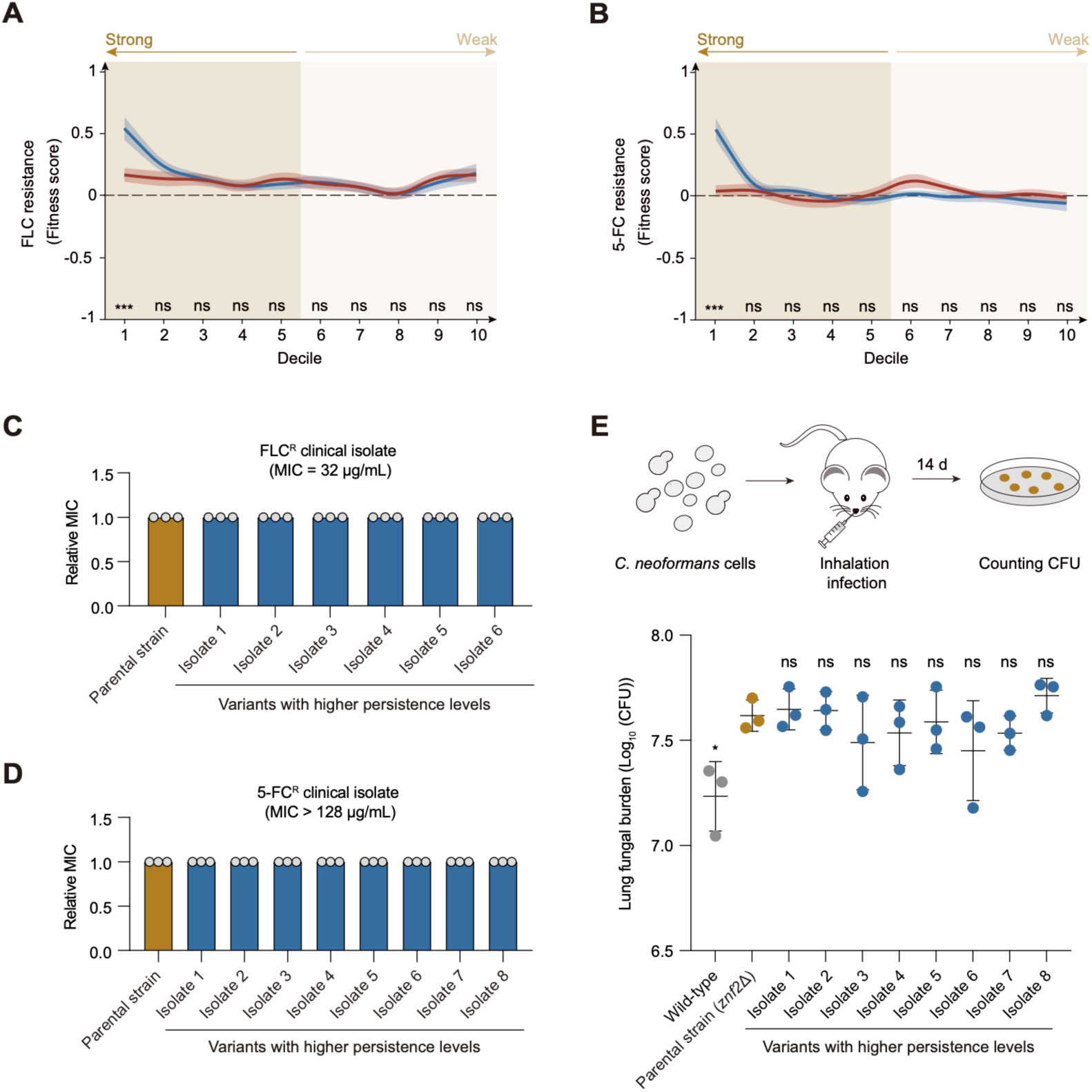
AmB persistence evolution can occur in fungistatic-resistant and hypervirulent backgrounds, related to Figure 3. (A) Fitness scores under FLC exposure across deciles ranked by AmB resistance or AmB persistence. (B) Fitness scores under 5-FC exposure across deciles ranked by AmB resistance or AmB persistence. (C) Relative FLC MICs of variants with elevated persistence levels evolved from a FLC^R^ clinical isolate as the parental strain. Data shown as mean ± SD (n = 3). (D) Relative 5-FC MICs of variants with elevated persistence levels evolved from a 5-FC^R^ clinical isolate as the parental strain. Data shown as mean ± SD (n = 3). (E) Lung fungal burdens in mice infected with H99 wild-type, parental *znf2*Δ strain, and variants with elevated persistence levels evolved from the *znf2*Δ background. Data shown as mean ± SD (n = 3). ns, not significant; \**p* < 0.05; \*\*\**p* < 0.001.

**Figure S4.**
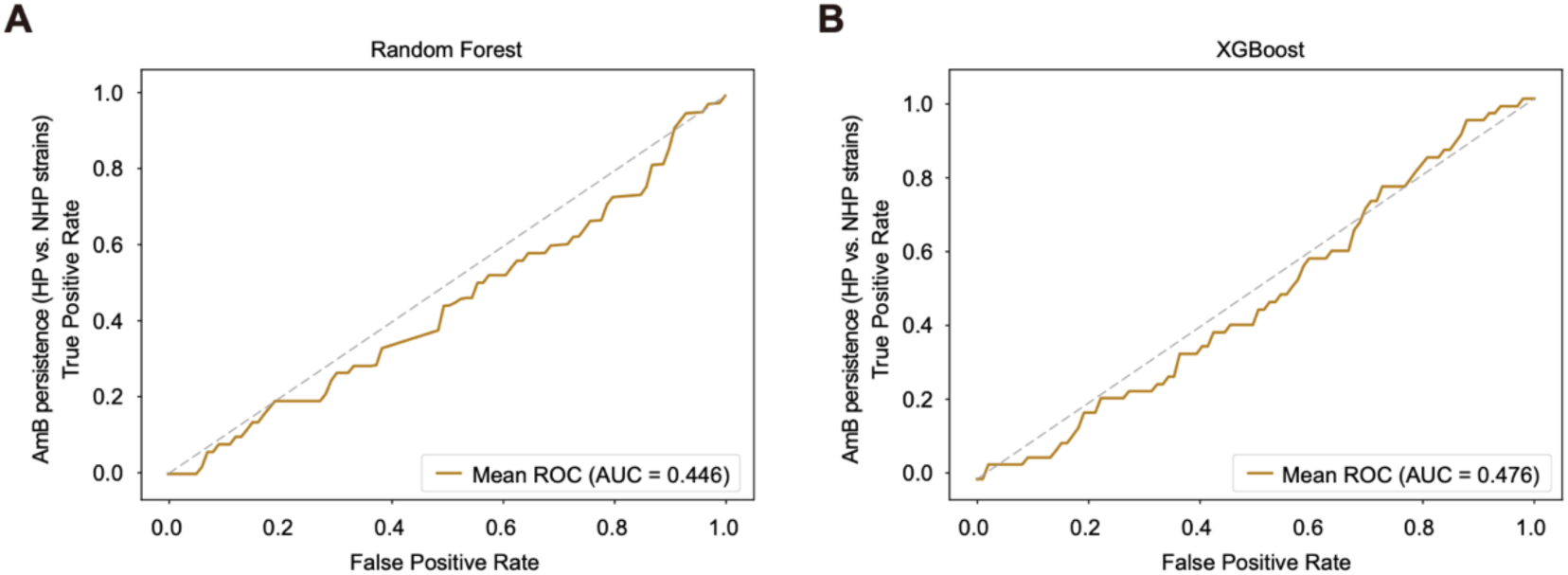
Genome-wide genetic variants poorly predict HP clinical isolates, related to Figure 4. (A) ROC curve for predicting HP clinical isolates using a Random Forest model based on genome-wide variants. (B) ROC curve for predicting HP clinical isolates using a XGBoost model based on genome-wide variants.

**Figure S5.**
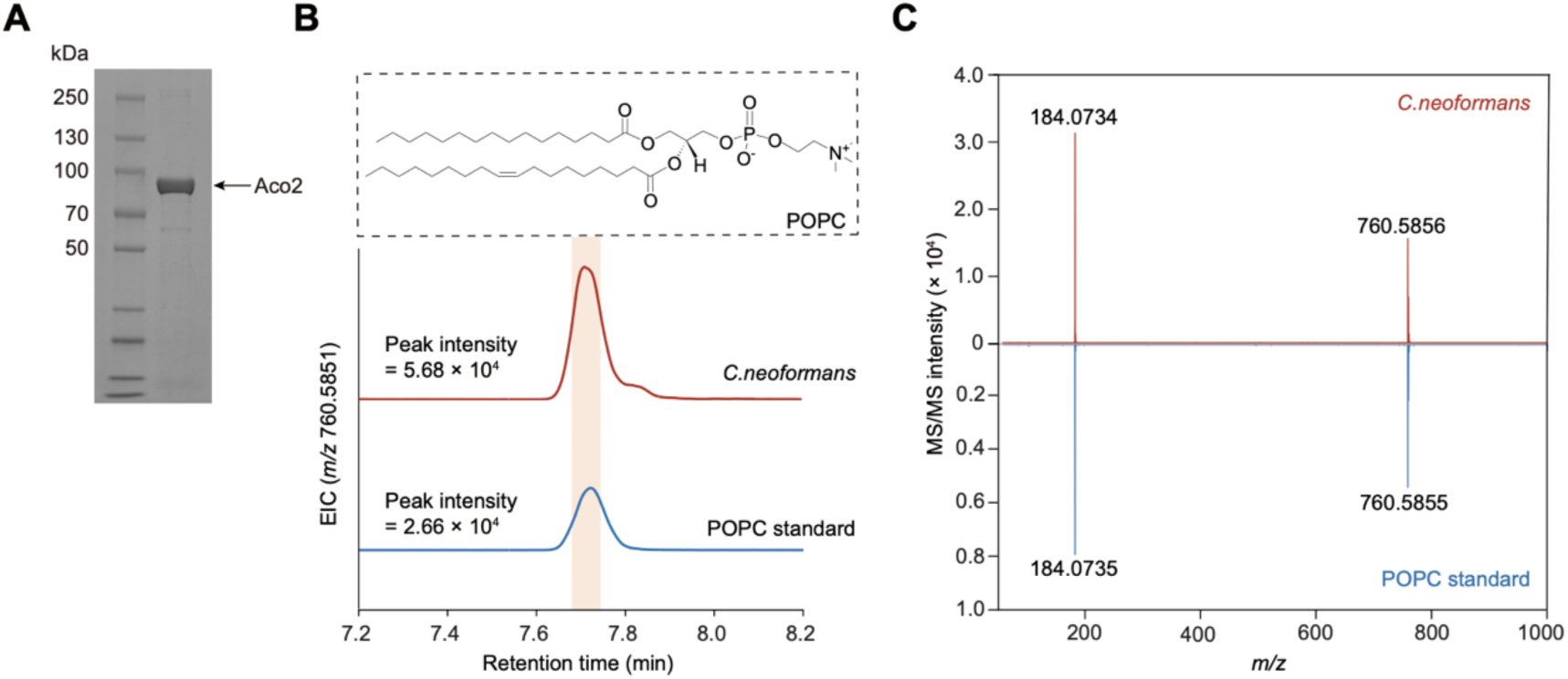
Validation of recombinant Aco2 purification and POPC detection, related to Figure 5. (A) Coomassie brilliant blue-stained SDS-PAGE showing purified Aco2 (shown in grayscale). (B) EIC at *m/z* 760.5851 ([M+H]^+^) from LC-MS analysis of the *C.neoformans* SPCs extract and POPC standard. (C) Tandem mass spectrometry (MS/MS) analysis of the corresponding peaks shown in (C). *m/z* ([M+H]^+^) values for the two major peaks in each trace are annotated. The characteristic phosphocholine fragment at *m/z* 184.073 was detected in both spectra.^156^

## Notes

### Competing Interest Statement

The authors have declared no competing interest.

## REFERENCES

1. Denning, D.W. (2024). Global incidence and mortality of severe fungal disease. Lancet Infect. Dis. 24, e428–e438. 10.1016/S1473-3099(23)00692-8.

2. Fisher, M.C., A. Alastruey-Izquierdo, J. Berman, T. Bicanic, E.M. Bignell, P. Bowyer, M. Bromley, R. Bruggemann, G. Garber, O.A. Cornely, et al. (2022). Tackling the emerging threat of antifungal resistance to human health. Nat. Rev. Microbiol. 20, 557–571. 10.1038/s41579-022-00720-1.

3. van Rhijn, N. and J. Rhodes (2025). Evolution of antifungal resistance in the environment. Nat Microbiol 10, 1804–1815. 10.1038/s41564-025-02055-y.

4. Carmona, E.M. and A.H. Limper (2017). Overview of Treatment Approaches for Fungal Infections. Clin. Chest Med. 38, 393–402. 10.1016/j.ccm.2017.04.003.

5. Reddy, G.K.K., A.R. Padmavathi, and Y.V. Nancharaiah (2022). Fungal infections: Pathogenesis, antifungals and alternate treatment approaches. Curr Res Microb Sci 3, 100137. 10.1016/j.crmicr.2022.100137.

6. Sun, S., M.J. Hoy, and J. Heitman (2020). Fungal pathogens. Curr. Biol. 30, R1163–R1169. 10.1016/j.cub.2020.07.032.

7. Cavassin, F.B., J.L. Bau-Carneiro, R.R. Vilas-Boas, and F. Queiroz-Telles (2021). Sixty years of Amphotericin B: An Overview of the Main Antifungal Agent Used to Treat Invasive Fungal Infections. Infect Dis Ther 10, 115–147. 10.1007/s40121-020-00382-7.

8. Chang, C.C., T.S. Harrison, T.A. Bicanic, M. Chayakulkeeree, T.C. Sorrell, A. Warris, F. Hagen, A. Spec, R. Oladele, N.P. Govender, et al. (2024). Global guideline for the diagnosis and management of cryptococcosis: an initiative of the ECMM and ISHAM in cooperation with the ASM. Lancet Infect. Dis. 24, e495–e512. 10.1016/S1473-3099(23)00731-4.

9. Iyer, K.R., N.M. Revie, C. Fu, N. Robbins, and L.E. Cowen (2021). Treatment strategies for cryptococcal infection: challenges, advances and future outlook. Nat. Rev. Microbiol. 19, 454–466. 10.1038/s41579-021-00511-0.

10. Williamson, P.R., J.N. Jarvis, A.A. Panackal, M.C. Fisher, S.F. Molloy, A. Loyse, and T.S. Harrison (2017). Cryptococcal meningitis: epidemiology, immunology, diagnosis and therapy. Nat. Rev. Neurol. 13, 13–24. 10.1038/nrneurol.2016.167.

11. May, R.C., N.R. Stone, D.L. Wiesner, T. Bicanic, and K. Nielsen (2016). Cryptococcus: from environmental saprophyte to global pathogen. Nat. Rev. Microbiol. 14, 106–17. 10.1038/nrmicro.2015.6.

12. Lin, X. and J. Heitman (2006). The biology of the Cryptococcus neoformans species complex. Annu. Rev. Microbiol. 60, 69–105. 10.1146/annurev.micro.60.080805.142102.

13. Bahn, Y.S., S. Sun, J. Heitman, and X. Lin (2020). Microbe Profile: Cryptococcus neoformans species complex. Microbiology (Reading) 166, 797–799. 10.1099/mic.0.000973.

14. Brown, J.C.S., J. Nelson, B. VanderSluis, R. Deshpande, A. Butts, S. Kagan, I. Polacheck, D.J. Krysan, C.L. Myers, and H.D. Madhani (2014). Unraveling the biology of a fungal meningitis pathogen using chemical genetics. Cell 159, 1168–1187. 10.1016/j.cell.2014.10.044.

15. Tugume, L., K. Ssebambulidde, J. Kasibante, J. Ellis, R.M. Wake, J. Gakuru, D.S. Lawrence, M. Abassi, R. Rajasingham, D.B. Meya, et al. (2023). Cryptococcal meningitis. Nat Rev Dis Primers 9, 62. 10.1038/s41572-023-00472-z.

16. Fisher, M.C. and D.W. Denning (2023). The WHO fungal priority pathogens list as a game-changer. Nat. Rev. Microbiol. 21, 211–212. 10.1038/s41579-023-00861-x.

17. Rajasingham, R., N.P. Govender, A. Jordan, A. Loyse, A. Shroufi, D.W. Denning, D.B. Meya, T.M. Chiller, and D.R. Boulware (2022). The global burden of HIV-associated cryptococcal infection in adults in 2020: a modelling analysis. Lancet Infect. Dis. 22, 1748–1755. 10.1016/S1473-3099(22)00499-6.

18. Zhao, Y., L. Ye, F. Zhao, L. Zhang, Z. Lu, T. Chu, S. Wang, Z. Liu, Y. Sun, M. Chen, et al. (2023). Cryptococcus neoformans, a global threat to human health. Infect Dis Poverty 12, 20. 10.1186/s40249-023-01073-4.

19. Caballero-Flores, G., J.M. Pickard, and G. Nunez (2023). Microbiota-mediated colonization resistance: mechanisms and regulation. Nat. Rev. Microbiol. 21, 347–360. 10.1038/s41579-022-00833-7.

20. Brauner, A., O. Fridman, O. Gefen, and N.Q. Balaban (2016). Distinguishing between resistance, tolerance and persistence to antibiotic treatment. Nat. Rev. Microbiol. 14, 320–30. 10.1038/nrmicro.2016.34.

21. Espinel-Ingroff, A., A. Chowdhary, M. Cuenca-Estrella, A. Fothergill, J. Fuller, F. Hagen, N. Govender, J. Guarro, E. Johnson, C. Lass-Florl, et al. (2012). Cryptococcus neoformans-Cryptococcus gattii species complex: an international study of wild-type susceptibility endpoint distributions and epidemiological cutoff values for amphotericin B and flucytosine. Antimicrob. Agents Chemother. 56, 3107–13. 10.1128/AAC.06252-11.

22. Davey, K.G., A.D. Holmes, E.M. Johnson, A. Szekely, and D.W. Warnock (1998). Comparative evaluation of FUNGITEST and broth microdilution methods for antifungal drug susceptibility testing of Candida species and Cryptococcus neoformans. J. Clin. Microbiol. 36, 926–30. 10.1128/JCM.36.4.926-930.1998.

23. Smith, K.D., B. Achan, K.H. Hullsiek, T.R. McDonald, L.H. Okagaki, A.A. Alhadab, A. Akampurira, J.R. Rhein, D.B. Meya, D.R. Boulware, et al. (2015). Increased Antifungal Drug Resistance in Clinical Isolates of Cryptococcus neoformans in Uganda. Antimicrob. Agents Chemother. 59, 7197–204. 10.1128/AAC.01299-15.

24. Pfaller, M.A., S.A. Messer, L. Boyken, C. Rice, S. Tendolkar, R.J. Hollis, G.V. Doern, and D.J. Diekema (2005). Global trends in the antifungal susceptibility of Cryptococcus neoformans (1990 to 2004). J. Clin. Microbiol. 43, 2163–7. 10.1128/JCM.43.5.2163-2167.2005.

25. Pfaller, M.A. and D.J. Diekema (2012). Progress in antifungal susceptibility testing of Candida spp. by use of Clinical and Laboratory Standards Institute broth microdilution methods, 2010 to 2012. J. Clin. Microbiol. 50, 2846–56. 10.1128/JCM.00937-12.

26. Ahmad, S., L. Joseph, J.E. Parker, M. Asadzadeh, S.L. Kelly, J.F. Meis, and Z. Khan (2019). ERG6 and ERG2 Are Major Targets Conferring Reduced Susceptibility to Amphotericin B in Clinical Candida glabrata Isolates in Kuwait. Antimicrob. Agents Chemother. 63, 10.1128/AAC.01900-18.

27. Vincent, B.M., A.K. Lancaster, R. Scherz-Shouval, L. Whitesell, and S. Lindquist (2013). Fitness trade-offs restrict the evolution of resistance to amphotericin B. PLoS Biol. 11, e1001692. 10.1371/journal.pbio.1001692.

28. Lee, Y., N. Robbins, and L.E. Cowen (2023). Molecular mechanisms governing antifungal drug resistance. NPJ Antimicrob Resist 1, 5. 10.1038/s44259-023-00007-2.

29. Rolfes, M.A., J. Rhein, C. Schutz, K. Taseera, H.W. Nabeta, K. Huppler Hullsiek, A. Akampuira, R. Rajasingham, A. Musubire, D.A. Williams, et al. (2015). Cerebrospinal Fluid Culture Positivity and Clinical Outcomes After Amphotericin-Based Induction Therapy for Cryptococcal Meningitis. Open Forum Infect Dis 2, ofv157. 10.1093/ofid/ofv157.

30. Bahr, N.C., C.P. Skipper, K. Huppler-Hullsiek, K. Ssebambulidde, B.M. Morawski, N.W. Engen, E. Nuwagira, C.M. Quinn, P.S. Ramachandran, E.E. Evans, et al. (2023). Recurrence of Symptoms Following Cryptococcal Meningitis: Characterizing a Diagnostic Conundrum With Multiple Etiologies. Clin. Infect. Dis. 76, 1080–1087. 10.1093/cid/ciac853.

31. Musubire, A.K., D.R. Boulware, D.B. Meya, and J. Rhein (2013). Diagnosis and Management of Cryptococcal Relapse. J. AIDS Clin. Res. Suppl 3, 10.4172/2155-6113.s3-003.

32. Zafar, H., S. Altamirano, E.R. Ballou, and K. Nielsen (2019). A titanic drug resistance threat in Cryptococcus neoformans. Curr. Opin. Microbiol. 52, 158–164. 10.1016/j.mib.2019.11.001.

33. Chen, L., L. Zhang, Y. Xie, Y. Wang, X. Tian, W. Fang, X. Xue, and L. Wang (2023). Confronting antifungal resistance, tolerance, and persistence: Advances in drug target discovery and delivery systems. Adv Drug Deliv Rev 200, 115007. 10.1016/j.addr.2023.115007.

34. Ma, X., J. Cui, Y. Tao, G. Liao, and L. Wang (2026). Emergence of traits in human fungal pathogens. Trends Microbiol. 34, 497–508. 10.1016/j.tim.2025.12.005.

35. Berman, J. and D.J. Krysan (2020). Drug resistance and tolerance in fungi. Nat. Rev. Microbiol. 18, 319–331. 10.1038/s41579-019-0322-2.

36. Chen, L., X. Tian, L. Zhang, W. Wang, P. Hu, Z. Ma, Y. Li, S. Li, Z. Shen, X. Fan, et al. (2024). Brain glucose induces tolerance of Cryptococcus neoformans to amphotericin B during meningitis. Nat Microbiol 9, 346–358. 10.1038/s41564-023-01561-1.

37. Yang, F. and J. Berman (2024). Beyond resistance: antifungal heteroresistance and antifungal tolerance in fungal pathogens. Curr. Opin. Microbiol. 78, 102439. 10.1016/j.mib.2024.102439.

38. Bojsen, R., B. Regenberg, and A. Folkesson (2017). Persistence and drug tolerance in pathogenic yeast. Curr. Genet. 63, 19–22. 10.1007/s00294-016-0613-3.

39. Ronneau, S., P.W. Hill, and S. Helaine (2021). Antibiotic persistence and tolerance: not just one and the same. Curr. Opin. Microbiol. 64, 76–81. 10.1016/j.mib.2021.09.017.

40. Lewis, K. (2010). Persister cells. Annu. Rev. Microbiol. 64, 357–72. 10.1146/annurev.micro.112408.134306.

41. Balaban, N.Q., S. Helaine, K. Lewis, M. Ackermann, B. Aldridge, D.I. Andersson, M.P. Brynildsen, D. Bumann, A. Camilli, J.J. Collins, et al. (2019). Definitions and guidelines for research on antibiotic persistence. Nat. Rev. Microbiol. 17, 441–448. 10.1038/s41579-019-0196-3.

42. Xie, Y., W. Ke, K.H. Wong, and L. Wang (2025). Persister cells in human fungal pathogens. PLoS Pathog. 21, e1013483. 10.1371/journal.ppat.1013483.

43. Fisher, R.A., B. Gollan, and S. Helaine (2017). Persistent bacterial infections and persister cells. Nat. Rev. Microbiol. 15, 453–464. 10.1038/nrmicro.2017.42.

44. Monack, D.M., A. Mueller, and S. Falkow (2004). Persistent bacterial infections: the interface of the pathogen and the host immune system. Nat. Rev. Microbiol. 2, 747–65. 10.1038/nrmicro955.

45. Cabral, D.J., J.I. Wurster, and P. Belenky (2018). Antibiotic Persistence as a Metabolic Adaptation: Stress, Metabolism, the Host, and New Directions. Pharmaceuticals (Basel) 11, 10.3390/ph11010014.

46. Gollan, B., G. Grabe, C. Michaux, and S. Helaine (2019). Bacterial Persisters and Infection: Past, Present, and Progressing. Annu. Rev. Microbiol. 73, 359–385. 10.1146/annurev-micro-020518-115650.

47. Bakkeren, E., M. Diard, and W.D. Hardt (2020). Evolutionary causes and consequences of bacterial antibiotic persistence. Nat. Rev. Microbiol. 18, 479–490. 10.1038/s41579-020-0378-z.

48. Brown, J.C. and E.R. Ballou (2024). Is Cryptococcus neoformans a pleomorphic fungus? Curr. Opin. Microbiol. 82, 102539. 10.1016/j.mib.2024.102539.

49. Maisonneuve, E. and K. Gerdes (2014). Molecular mechanisms underlying bacterial persisters. Cell 157, 539–48. 10.1016/j.cell.2014.02.050.

50. Claudi, B., P. Sprote, A. Chirkova, N. Personnic, J. Zankl, N. Schurmann, A. Schmidt, and D. Bumann (2014). Phenotypic variation of Salmonella in host tissues delays eradication by antimicrobial chemotherapy. Cell 158, 722–733. 10.1016/j.cell.2014.06.045.

51. Al-Dhaheri, R.S. and L.J. Douglas (2008). Absence of amphotericin B-tolerant persister cells in biofilms of some Candida species. Antimicrob. Agents Chemother. 52, 1884–7. 10.1128/AAC.01473-07.

52. Arastehfar, A., F. Daneshnia, D.J. Floyd, N.E. Jeffries, M. Salehi, D.S. Perlin, M. Ilkit, C. Lass-Floerl, and M.K. Mansour (2024). Echinocandin persistence directly impacts the evolution of resistance and survival of the pathogenic fungus Candida glabrata. mBio 15, e0007224. 10.1128/mbio.00072-24.

53. Harrison, J.J., R.J. Turner, and H. Ceri (2007). A subpopulation of Candida albicans and Candida tropicalis biofilm cells are highly tolerant to chelating agents. FEMS Microbiol. Lett. 272, 172–81. 10.1111/j.1574-6968.2007.00745.x.

54. Arastehfar, A., F. Daneshnia, N. Cabrera, S. Penalva-Lopez, J. Sarathy, M. Zimmerman, E. Shor, and D.S. Perlin (2023). Macrophage internalization creates a multidrug-tolerant fungal persister reservoir and facilitates the emergence of drug resistance. Nat Commun 14, 1183. 10.1038/s41467-023-36882-6.

55. LaFleur, M.D., C.A. Kumamoto, and K. Lewis (2006). Candida albicans biofilms produce antifungal-tolerant persister cells. Antimicrob. Agents Chemother. 50, 3839–46. 10.1128/AAC.00684-06.

56. Scott, J., C. Valero, A. Mato-Lopez, I.J. Donaldson, A. Roldan, H. Chown, N. Van Rhijn, R. Lobo-Vega, S. Gago, T. Furukawa, et al. (2023). Aspergillus fumigatus Can Display Persistence to the Fungicidal Drug Voriconazole. Microbiol Spectr 11, e0477022. 10.1128/spectrum.04770-22.

57. Ke, W., Y. Xie, Y. Chen, H. Ding, L. Ye, H. Qiu, H. Li, L. Zhang, L. Chen, X. Tian, et al. (2024). Fungicide-tolerant persister formation during cryptococcal pulmonary infection. Cell Host Microbe 32, 276–289 e7. 10.1016/j.chom.2023.12.012.

58. Sun, J., Z. Li, H. Chu, J. Guo, G. Jiang, and Q. Qi (2016). Candida albicans Amphotericin B-Tolerant Persister Formation is Closely Related to Surface Adhesion. Mycopathologia 181, 41–9. 10.1007/s11046-015-9894-1.

59. 59. (1997). Case definitions for infectious conditions under public health surveillance. Centers for Disease Control and Prevention. MMWR Recomm. Rep. 46, 1–55.

60. Miller, J.M., M.J. Binnicker, S. Campbell, K.C. Carroll, K.C. Chapin, M.D. Gonzalez, A. Harrington, R.C. Jerris, S.C. Kehl, S.M. Leal, Jr.,et al. (2024). Guide to Utilization of the Microbiology Laboratory for Diagnosis of Infectious Diseases: 2024 Update by the Infectious Diseases Society of America (IDSA) and the American Society for Microbiology (ASM). Clin. Infect. Dis. 10.1093/cid/ciae104.

61. Bojsen, R., B. Regenberg, D. Gresham, and A. Folkesson (2016). A common mechanism involving the TORC1 pathway can lead to amphotericin B-persistence in biofilm and planktonic Saccharomyces cerevisiae populations. Sci. Rep. 6, 21874. 10.1038/srep21874.

62. Alanio, A., F. Vernel-Pauillac, A. Sturny-Leclere, and F. Dromer (2015). Cryptococcus neoformans host adaptation: toward biological evidence of dormancy. mBio 6, 10.1128/mBio.02580-14.

63. Bojsen, R., B. Regenberg, and A. Folkesson (2014). Saccharomyces cerevisiae biofilm tolerance towards systemic antifungals depends on growth phase. BMC Microbiol. 14, 305. 10.1186/s12866-014-0305-4.

64. Arendrup, M.C., J. Guinea, S. Arikan-Akdagli, E.F.J. Meijer, J.F. Meis, J.B. Buil, E. Dannaoui, C.G. Giske, P. Lyskova, J. Meletiadis, et al. (2026). How to interpret MICs of amphotericin B, echinocandins and flucytosine against Candida auris (Candidozyma auris) according to the newly established European Committee for Antimicrobial Susceptibility Testing (EUCAST) breakpoints. Clin. Microbiol. Infect. 32, 56–61. 10.1016/j.cmi.2025.07.002.

65. Zuiderveld, K.J. (1994). Contrast Limited Adaptive Histogram Equalization. in Graphics gems, pp. 474–485

66. Polesel, A., G. Ramponi, and V.J. Mathews (2000). Image enhancement via adaptive unsharp masking. IEEE Trans Image Process 9, 505–10. 10.1109/83.826787.

67. Jiang, H., Q. Guo, X. Zhi, H. Li, and Y. Chen (2026). A weakly supervised framework for automated biological assay assessment. Virus Res. 363, 199677. 10.1016/j.virusres.2025.199677.

68. Schneider, C.A., W.S. Rasband, and K.W. Eliceiri (2012). NIH Image to ImageJ: 25 years of image analysis. Nat Methods 9, 671–5. 10.1038/nmeth.2089.

69. Zhang, J., C. Li, M.M. Rahaman, Y. Yao, P. Ma, J. Zhang, X. Zhao, T. Jiang, and M. Grzegorzek (2022). A comprehensive review of image analysis methods for microorganism counting: from classical image processing to deep learning approaches. Artif Intell Rev 55, 2875–2944. 10.1007/s10462-021-10082-4.

70. Loftus, B.J., E. Fung, P. Roncaglia, D. Rowley, P. Amedeo, D. Bruno, J. Vamathevan, M. Miranda, I.J. Anderson, J.A. Fraser, et al. (2005). The genome of the basidiomycetous yeast and human pathogen Cryptococcus neoformans. Science 307, 1321–4. 10.1126/science.1103773.

71. Fan, X., L. Chen, M. Chen, N. Zhang, H. Chang, M. He, Z. Shen, L. Zhang, H. Ding, Y. Xie, et al. (2024). Pan-omics-based characterization and prediction of highly multidrug-adapted strains from an outbreak fungal species complex. Innovation (Camb) 5, 100681. 10.1016/j.xinn.2024.100681.

72. Fan, X., M. Xiao, S. Chen, F. Kong, H.T. Dou, H. Wang, Y.L. Xiao, M. Kang, Z.Y. Sun, Z.D. Hu, et al. (2016). Predominance of Cryptococcus neoformans var. grubii multilocus sequence type 5 and emergence of isolates with non-wild-type minimum inhibitory concentrations to fluconazole: a multi-centre study in China. Clin. Microbiol. Infect. 22, 887 e1–887 e9. 10.1016/j.cmi.2016.07.008.

73. Rhodes, J., M.A. Beale, M. Vanhove, J.N. Jarvis, S. Kannambath, J.A. Simpson, A. Ryan, G. Meintjes, T.S. Harrison, M.C. Fisher, et al. (2017). A Population Genomics Approach to Assessing the Genetic Basis of Within-Host Microevolution Underlying Recurrent Cryptococcal Meningitis Infection. G3 (Bethesda) 7, 1165–1176. 10.1534/g3.116.037499.

74. Chen, Y., R.A. Farrer, C. Giamberardino, S. Sakthikumar, A. Jones, T. Yang, J.L. Tenor, O. Wagih, M. Van Wyk, N.P. Govender, et al. (2017). Microevolution of Serial Clinical Isolates of Cryptococcus neoformans var. grubii and C. gattii. mBio 8, 10.1128/mBio.00166-17.

75. Ke, W., Y. Xie, Y. Hu, H. Ding, X. Fan, J. Huang, X. Tian, B. Zhang, Y. Xu, X. Liu, et al. (2022). A forkhead transcription factor contributes to the regulatory differences of pathogenicity in closely related fungal pathogens. mLife 1, 79–91. 10.1002/mlf2.12011.

76. Mukaremera, L., T.R. McDonald, J.N. Nielsen, C.J. Molenaar, A. Akampurira, C. Schutz, K. Taseera, C. Muzoora, G. Meintjes, D.B. Meya, et al. (2019). The Mouse Inhalation Model of Cryptococcus neoformans Infection Recapitulates Strain Virulence in Humans and Shows that Closely Related Strains Can Possess Differential Virulence. Infect. Immun. 87, 10.1128/IAI.00046-19.

77. Liu, O.W., C.D. Chun, E.D. Chow, C. Chen, H.D. Madhani, and S.M. Noble (2008). Systematic genetic analysis of virulence in the human fungal pathogen Cryptococcus neoformans. Cell 135, 174–88. 10.1016/j.cell.2008.07.046.

78. Oliveira, F.F.M., H.C. Paes, L.D.F. Peconick, F.L. Fonseca, C.L.F. Marina, A.L. Bocca, M. Homem-de-Mello, M.L. Rodrigues, P. Albuquerque, A.M. Nicola, et al. (2020). Erg6 affects membrane composition and virulence of the human fungal pathogen Cryptococcus neoformans. Fungal Genet. Biol. 140, 103368. 10.1016/j.fgb.2020.103368.

79. Boucher, M.J., S. Banerjee, M.B. Joshi, A.L. Wei, M.J. Nalley, M.Y. Huang, S. Lei, M. Ciranni, A. Condon, A. Langen, et al. (2025). Phenotypic landscape of an invasive fungal pathogen reveals its unique biology. Cell 188, 4003–4024 e24. 10.1016/j.cell.2025.05.017.

80. Wang, L., B. Zhai, and X. Lin (2012). The link between morphotype transition and virulence in Cryptococcus neoformans. PLoS Pathog. 8, e1002765. 10.1371/journal.ppat.1002765.

81. Arendrup, M.C., A. Prakash, J. Meletiadis, C. Sharma, and A. Chowdhary (2017). Comparison of EUCAST and CLSI Reference Microdilution MICs of Eight Antifungal Compounds for Candida auris and Associated Tentative Epidemiological Cutoff Values. Antimicrob. Agents Chemother. 61, 10.1128/AAC.00485-17.

82. Wang, T., W. Shao, Z. Huang, H. Tang, J. Zhang, Z. Ding, and K. Huang (2021). MOGONET integrates multi-omics data using graph convolutional networks allowing patient classification and biomarker identification. Nat Commun 12, 3445. 10.1038/s41467-021-23774-w.

83. Hedou, J., I. Maric, G. Bellan, J. Einhaus, D.K. Gaudilliere, F.X. Ladant, F. Verdonk, I.A. Stelzer, D. Feyaerts, A.S. Tsai, et al. (2024). Discovery of sparse, reliable omic biomarkers with Stabl. Nat. Biotechnol. 42, 1581–1593. 10.1038/s41587-023-02033-x.

84. Nguyen, P.B.H., D. Garger, D. Lu, H. Maalmi, H. Prokisch, B. Thorand, J. Adamski, G. Kastenmuller, M. Waldenberger, C. Gieger, et al. (2024). Interpretable multimodal machine learning (IMML) framework reveals pathological signatures of distal sensorimotor polyneuropathy. Commun Med (Lond) 4, 265. 10.1038/s43856-024-00637-1.

85. Zhang, C., C. Wang, Q.S. Hu, X.M. Xu, R.H. Yan, X.L. Nie, Y.G. Peng, H.P. Yang, Y. Song, X.J. Yang, et al. (2025). Development and validation of a real-time risk prediction model for acute kidney injury in hospitalized pediatric patients. World J. Pediatr. 21, 878–888. 10.1007/s12519-025-00950-2.

86. Moshkov, N., T. Becker, K. Yang, P. Horvath, V. Dancik, B.K. Wagner, P.A. Clemons, S. Singh, A.E. Carpenter, and J.C. Caicedo (2023). Predicting compound activity from phenotypic profiles and chemical structures. Nat Commun 14, 1967. 10.1038/s41467-023-37570-1.

87. Wang, P., M.D. Lehti-Shiu, S. Lotreck, K. Segura Aba, P.J. Krysan, and S.H. Shiu (2024). Prediction of plant complex traits via integration of multi-omics data. Nat Commun 15, 6856. 10.1038/s41467-024-50701-6.

88. Carli, F., P. Di Chiaro, M. Morelli, C. Arora, L. Bisceglia, N. De Oliveira Rosa, A. Cortesi, S. Franceschi, F. Lessi, A.L. Di Stefano, et al. (2025). Learning and actioning general principles of cancer cell drug sensitivity. Nat Commun 16, 1654. 10.1038/s41467-025-56827-5.

89. Basenko, E.Y., J.A. Pulman, A. Shanmugasundram, O.S. Harb, K. Crouch, D. Starns, S. Warrenfeltz, C. Aurrecoechea, C.J. Stoeckert, Jr., J.C. Kissinger, et al. (2018). FungiDB: An Integrated Bioinformatic Resource for Fungi and Oomycetes. J Fungi (Basel) 4, 10.3390/jof4010039.

90. Caza, M., G. Hu, M. Price, J.R. Perfect, and J.W. Kronstad (2016). The Zinc Finger Protein Mig1 Regulates Mitochondrial Function and Azole Drug Susceptibility in the Pathogenic Fungus Cryptococcus neoformans. mSphere 1, 10.1128/mSphere.00080-15.

91. Xie, A., J.S. Brunner, S. Chakraborty, A.M. Montero, A.E. Bridgeman, K.I. Paras, R. Cui, M. Fagoaga-Eugui, M. Komza, P.K. Arnold, et al. (2026). Citrate clearance is a major function of aconitase 2 in the canonical TCA cycle. Cell 189, 2684–2699 e21. 10.1016/j.cell.2026.01.028.

92. Lushchak, O.V., M. Piroddi, F. Galli, and V.I. Lushchak (2014). Aconitase post-translational modification as a key in linkage between Krebs cycle, iron homeostasis, redox signaling, and metabolism of reactive oxygen species. Redox Rep 19, 8–15. 10.1179/1351000213Y.0000000073.

93. Shi, R. and X. Lin (2024). Illuminating the Cryptococcus neoformans species complex: unveiling intracellular structures with fluorescent-protein-based markers. Genetics 227, 10.1093/genetics/iyae059.

94. Jung, S.J., Y. Seo, K.C. Lee, D. Lee, and J.H. Roe (2015). Essential function of Aco2, a fusion protein of aconitase and mitochondrial ribosomal protein bL21, in mitochondrial translation in fission yeast. FEBS Lett. 589, 822–8. 10.1016/j.febslet.2015.02.015.

95. Mansilla, S., V. Tortora, F. Pignataro, S. Sastre, I. Castro, M.L. Chiribao, C. Robello, A. Zeida, J. Santos, and L. Castro (2023). Redox sensitive human mitochondrial aconitase and its interaction with frataxin: In vitro and in silico studies confirm that it takes two to tango. Free Radic. Biol. Med. 197, 71–84. 10.1016/j.freeradbiomed.2023.01.028.

96. McQuaw, C.M., L. Zheng, A.G. Ewing, and N. Winograd (2007). Localization of sphingomyelin in cholesterol domains by imaging mass spectrometry. Langmuir 23, 5645–50. 10.1021/la063251f.

97. Zhang, H., Y. Liu, L. Fields, X. Shi, P. Huang, H. Lu, A.J. Schneider, X. Tang, L. Puglielli, N.V. Welham, et al. (2023). Single-cell lipidomics enabled by dual-polarity ionization and ion mobility-mass spectrometry imaging. Nat Commun 14, 5185. 10.1038/s41467-023-40512-6.

98. Lewis, K. (2020). The Science of Antibiotic Discovery. Cell 181, 29–45. 10.1016/j.cell.2020.02.056.

99. Butts, A., L. DiDone, K. Koselny, B.K. Baxter, Y. Chabrier-Rosello, M. Wellington, and D.J. Krysan (2013). A repurposing approach identifies off-patent drugs with fungicidal cryptococcal activity, a common structural chemotype, and pharmacological properties relevant to the treatment of cryptococcosis. Eukaryot Cell 12, 278–87. 10.1128/EC.00314-12.

100. Organization, W.H., Antifungal agents in clinical and preclinical development: overview and analysis. 2025, World Health Organization. p. 88.

101. Mitsuyama, J., N. Nomura, K. Hashimoto, E. Yamada, H. Nishikawa, M. Kaeriyama, A. Kimura, Y. Todo, and H. Narita (2008). In vitro and in vivo antifungal activities of T-2307, a novel arylamidine. Antimicrob. Agents Chemother. 52, 1318–24. 10.1128/AAC.01159-07.

102. Nishikawa, H., Y. Fukuda, J. Mitsuyama, M. Tashiro, A. Tanaka, T. Takazono, T. Saijo, K. Yamamoto, S. Nakamura, Y. Imamura, et al. (2017). In vitro and in vivo antifungal activities of T-2307, a novel arylamidine, against Cryptococcus gattii: an emerging fungal pathogen. J. Antimicrob. Chemother. 72, 1709–1713. 10.1093/jac/dkx020.

103. Giamberardino, C.D., J.L. Tenor, D.L. Toffaletti, J.R. Palmucci, W. Schell, J.V. Boua, C. Marius, K.E. Stott, S.L. Steele, W. Hope, et al. (2023). Pharmacodynamics of ATI-2307 in a rabbit model of cryptococcal meningoencephalitis. Antimicrob. Agents Chemother. 67, e0081823. 10.1128/aac.00818-23.

104. Shibata, T., T. Takahashi, E. Yamada, A. Kimura, H. Nishikawa, H. Hayakawa, N. Nomura, and J. Mitsuyama (2012). T-2307 causes collapse of mitochondrial membrane potential in yeast. Antimicrob. Agents Chemother. 56, 5892–7. 10.1128/AAC.05954-11.

105. Wiederhold, N.P. (2021). Review of T-2307, an Investigational Agent That Causes Collapse of Fungal Mitochondrial Membrane Potential. J Fungi (Basel) 7, 10.3390/jof7020130.

106. McCarty, T.P. and P.G. Pappas (2021). Antifungal Pipeline. Front Cell Infect Microbiol 11, 732223. 10.3389/fcimb.2021.732223.

107. Lafleur, M.D., Q. Qi, and K. Lewis (2010). Patients with long-term oral carriage harbor high-persister mutants of Candida albicans. Antimicrob. Agents Chemother. 54, 39–44. 10.1128/AAC.00860-09.

108. Faigenbaum-Romm, R., N. Yedidi, O. Gefen, N. Katsowich-Nagar, L. Aroeti, I. Ronin, M. Bar-Meir, I. Rosenshine, and N.Q. Balaban (2025). Uncovering phenotypic inheritance from single cells with Microcolony-seq. Cell 188, 5313–5331 e18. 10.1016/j.cell.2025.08.001.

109. Ma, P., H.M. Amemiya, L.L. He, S.J. Gandhi, R. Nicol, R.P. Bhattacharyya, C.S. Smillie, and D.T. Hung (2023). Bacterial droplet-based single-cell RNA-seq reveals antibiotic-associated heterogeneous cellular states. Cell 186, 877–891 e14. 10.1016/j.cell.2023.01.002.

110. Ejsing, C.S., J.L. Sampaio, V. Surendranath, E. Duchoslav, K. Ekroos, R.W. Klemm, K. Simons, and A. Shevchenko (2009). Global analysis of the yeast lipidome by quantitative shotgun mass spectrometry. Proc. Natl. Acad. Sci. U. S. A. 106, 2136–41. 10.1073/pnas.0811700106.

111. Kuge, H., K. Akahori, K.I. Yagyu, and K. Honke (2014). Functional compartmentalization of the plasma membrane of neurons by a unique acyl chain composition of phospholipids. J. Biol. Chem. 289, 26783–26793. 10.1074/jbc.M114.571075.

112. Li, Z., S. Cheng, Q. Lin, W. Cao, J. Yang, M. Zhang, A. Shen, W. Zhang, Y. Xia, X. Ma, et al. (2021). Single-cell lipidomics with high structural specificity by mass spectrometry. Nat Commun 12, 2869. 10.1038/s41467-021-23161-5.

113. Mirel, D.B., K. Marder, J. Graziano, G. Freyer, Q. Zhao, R. Mayeux, and K.C. Wilhelmsen (1998). Characterization of the human mitochondrial aconitase gene (ACO2). Gene 213, 205–18. 10.1016/s0378-1119(98)00188-7.

114. Lorent, J.H., K.R. Levental, L. Ganesan, G. Rivera-Longsworth, E. Sezgin, M. Doktorova, E. Lyman, and I. Levental (2020). Plasma membranes are asymmetric in lipid unsaturation, packing and protein shape. Nat. Chem. Biol. 16, 644–652. 10.1038/s41589-020-0529-6.

115. Cox, J.V., Y.M. Abdelrahman, J. Peters, N. Naher, and R.J. Belland (2016). Chlamydia trachomatis utilizes the mammalian CLA1 lipid transporter to acquire host phosphatidylcholine essential for growth. Cell. Microbiol. 18, 305–18. 10.1111/cmi.12523.

116. Itoe, M.A., J.L. Sampaio, G.G. Cabal, E. Real, V. Zuzarte-Luis, S. March, S.N. Bhatia, F. Frischknecht, C. Thiele, A. Shevchenko, et al. (2014). Host cell phosphatidylcholine is a key mediator of malaria parasite survival during liver stage infection. Cell Host Microbe 16, 778–86. 10.1016/j.chom.2014.11.006.

117. Lin, M., G. Grandinetti, L.M. Hartnell, D. Bliss, S. Subramaniam, and Y. Rikihisa (2020). Host membrane lipids are trafficked to membranes of intravacuolar bacterium Ehrlichia chaffeensis. Proc. Natl. Acad. Sci. U. S. A. 117, 8032–8043. 10.1073/pnas.1921619117.

118. Lozano-Chiu, M., V.L. Paetznick, M.A. Ghannoum, and J.H. Rex (1998). Detection of resistance to amphotericin B among Cryptococcus neoformans clinical isolates: performances of three different media assessed by using E-test and National Committee for Clinical Laboratory Standards M27-A methodologies. J. Clin. Microbiol. 36, 2817–22. 10.1128/JCM.36.10.2817-2822.1998.

119. Lin, J., Y. Fan, and X. Lin (2020). Transformation of Cryptococcus neoformans by electroporation using a transient CRISPR-Cas9 expression (TRACE) system. Fungal Genet. Biol. 138, 103364. 10.1016/j.fgb.2020.103364.

120. Upadhya, R., W.C. Lam, B.T. Maybruck, M.J. Donlin, A.L. Chang, S. Kayode, K.L. Ormerod, J.A. Fraser, T.L. Doering, and J.K. Lodge (2017). A fluorogenic C. neoformans reporter strain with a robust expression of m-cherry expressed from a safe haven site in the genome. Fungal Genet. Biol. 108, 13–25. 10.1016/j.fgb.2017.08.008.

121. Arras, S.D., J.L. Chitty, K.L. Blake, B.L. Schulz, and J.A. Fraser (2015). A genomic safe haven for mutant complementation in Cryptococcus neoformans. PLoS One 10, e0122916. 10.1371/journal.pone.0122916.

122. Hope, M.J., M.B. Bally, G. Webb, and P.R. Cullis (1985). Production of large unilamellar vesicles by a rapid extrusion procedure: characterization of size distribution, trapped volume and ability to maintain a membrane potential. Biochim. Biophys. Acta 812, 55–65. 10.1016/0005-2736(85)90521-8.

123. Mayer, L.D., M.J. Hope, and P.R. Cullis (1986). Vesicles of variable sizes produced by a rapid extrusion procedure. Biochim. Biophys. Acta 858, 161–8. 10.1016/0005-2736(86)90302-0.

124. Beier, S., A.M. Bolger, M.E. Bolger, R. Schwacke, and B. Usadel (2026). Trimmomatic: a decade of feature-rich, high-performance NGS read preprocessing. Bioinformatics 42, 10.1093/bioinformatics/btag331.

125. Prjibelski, A., D. Antipov, D. Meleshko, A. Lapidus, and A. Korobeynikov (2020). Using SPAdes De Novo Assembler. Curr Protoc Bioinformatics 70, e102. 10.1002/cpbi.102.

126. Gurevich, A., V. Saveliev, N. Vyahhi, and G. Tesler (2013). QUAST: quality assessment tool for genome assemblies. Bioinformatics 29, 1072–5. 10.1093/bioinformatics/btt086.

127. Li, H. and R. Durbin (2010). Fast and accurate long-read alignment with Burrows-Wheeler transform. Bioinformatics 26, 589–95. 10.1093/bioinformatics/btp698.

128. Danecek, P., J.K. Bonfield, J. Liddle, J. Marshall, V. Ohan, M.O. Pollard, A. Whitwham, T. Keane, S.A. McCarthy, R.M. Davies, et al. (2021). Twelve years of SAMtools and BCFtools. Gigascience 10, 10.1093/gigascience/giab008.

129. DePristo, M.A., E. Banks, R. Poplin, K.V. Garimella, J.R. Maguire, C. Hartl, A.A. Philippakis, G. del Angel, M.A. Rivas, M. Hanna, et al. (2011). A framework for variation discovery and genotyping using next-generation DNA sequencing data. Nat. Genet. 43, 491–8. 10.1038/ng.806.

130. Meyer, W., D.M. Aanensen, T. Boekhout, M. Cogliati, M.R. Diaz, M.C. Esposto, M. Fisher, F. Gilgado, F. Hagen, S. Kaocharoen, et al. (2009). Consensus multi-locus sequence typing scheme for Cryptococcus neoformans and Cryptococcus gattii. Med. Mycol. 47, 561–70. 10.1080/13693780902953886.

131. Ortiz, E.M. (2019). vcf2phylip v2.0: convert a VCF matrix into several matrix formats for phylogenetic analysis. Zenodo. 10.5281/zenodo.2540861.

132. Wong, T.K.F., N. Ly-Trong, H. Ren, P. Demotte, H. Banos, A.J. Roger, E. Susko, C. Bielow, N. De Maio, N. Goldman, et al. (2026). IQ-TREE 3: phylogenomic inference software using complex evolutionary models. Mol. Biol. Evol. 43, 10.1093/molbev/msag117.

133. Fan, X., R.C. Dai, S. Zhang, Y.Y. Geng, M. Kang, D.W. Guo, Y.N. Mei, Y.H. Pan, Z.Y. Sun, Y.C. Xu, et al. (2024). Author Correction: Tandem gene duplications contributed to high-level azole resistance in a rapidly expanding Candida tropicalis population. Nat Commun 15, 587. 10.1038/s41467-024-44825-y.

134. Alexander, D.H., J. Novembre, and K. Lange (2009). Fast model-based estimation of ancestry in unrelated individuals. Genome Res. 19, 1655–64. 10.1101/gr.094052.109.

135. Purcell, S., B. Neale, K. Todd-Brown, L. Thomas, M.A. Ferreira, D. Bender, J. Maller, P. Sklar, P.I. de Bakker, M.J. Daly, et al. (2007). PLINK: a tool set for whole-genome association and population-based linkage analyses. Am. J. Hum. Genet. 81, 559–75. 10.1086/519795.

136. Singh, A., A. MacKenzie, G. Girnun, and M. Del Poeta (2017). Analysis of sphingolipids, sterols, and phospholipids in human pathogenic Cryptococcus strains. J. Lipid Res. 58, 2017–2036. 10.1194/jlr.M078600.

137. Sun, X., W. Wang, K. Wang, X. Yu, J. Liu, F. Zhou, B. Xie, and S. Li (2013). Sterol C-22 Desaturase ERG5 Mediates the Sensitivity to Antifungal Azoles in Neurospora crassa and Fusarium verticillioides. Front. Microbiol. 4, 127. 10.3389/fmicb.2013.00127.

138. Matyash, V., G. Liebisch, T.V. Kurzchalia, A. Shevchenko, and D. Schwudke (2008). Lipid extraction by methyl-tert-butyl ether for high-throughput lipidomics. J. Lipid Res. 49, 1137–46. 10.1194/jlr.D700041-JLR200.

139. Vrkoslav, V., K. Prazakova, S. Strnad, K. Paukner, B. Muffova, S. Kauerova, J. Fronek, D. Sykora, J. Cvacka, R. Poledne, et al. (2025). Lipidomics of polarized macrophages in the human adipose tissue. Sci. Rep. 16, 3018. 10.1038/s41598-025-32912-z.

140. Wu, J., H. Jiang, Q. Bi, Q. Luo, J. Li, Y. Zhang, Z. Chen, and C. Li (2014). Apamin-mediated actively targeted drug delivery for treatment of spinal cord injury: more than just a concept. Mol. Pharm. 11, 3210–22. 10.1021/mp500393m.

141. Bernhard, M., N. Worasilchai, M. Kangogo, C. Bii, W.J. Trzaska, M. Weig, U. Gross, A. Chindamporn, and O. Bader (2021). CryptoType - Public Datasets for MALDI-TOF-MS Based Differentiation of Cryptococcus neoformans/gattii Complexes. Front Cell Infect Microbiol 11, 634382. 10.3389/fcimb.2021.634382.

142. Weis, C., A. Cuenod, B. Rieck, O. Dubuis, S. Graf, C. Lang, M. Oberle, M. Brackmann, K.K. Sogaard, M. Osthoff, et al. (2022). Direct antimicrobial resistance prediction from clinical MALDI-TOF mass spectra using machine learning. Nat. Med. 28, 164–174. 10.1038/s41591-021-01619-9.

143. Gibb, S. and K. Strimmer (2012). MALDIquant: a versatile R package for the analysis of mass spectrometry data. Bioinformatics 28, 2270–1. 10.1093/bioinformatics/bts447.

144. Wang, X., S. Wang, Z. Diao, X. Hou, Y. Gong, Q. Sun, J. Zhang, L. Ren, Y. Li, Y. Ji, et al. (2025). Label-free high-throughput live-cell sorting of genome-wide random mutagenesis libraries for metabolic traits by Raman flow cytometry. Proc. Natl. Acad. Sci. U. S. A. 122, e2503641122. 10.1073/pnas.2503641122.

145. Zhang, Y., G. Jing, R. Chen, Y. Gong, Y. Li, Y. Wang, X. Wang, J. Zhang, Y. Mao, Y. He, et al. (2026). RamEx: an R package for high-throughput microbial ramanome analyses with accurate quality assessment. Microbiome 14, 10.1186/s40168-026-02339-3.

146. Pedregosa, F., G. Varoquaux, A. Gramfort, V. Michel, B. Thirion, O. Grisel, M. Blondel, P. Prettenhofer, R. Weiss, V. Dubourg, et al. (2011). Scikit-learn: Machine Learning in Python. J. Mach. Learn. Res. 12, 2825–2830.

147. Chen, T. and C. Guestrin. (2016). XGBoost: A Scalable Tree Boosting System. Proceedings of the 22nd ACM SIGKDD International Conference on Knowledge Discovery and Data Mining. Association for Computing Machinery, pp. 785–794.

148. Lundberg, S.M. and S.-I. Lee. (2017). A unified approach to interpreting model predictions. Proceedings of the 31st International Conference on Neural Information Processing Systems, pp. 4768–4777.

149. Browning, B.L., Y. Zhou, and S.R. Browning (2018). A One-Penny Imputed Genome from Next-Generation Reference Panels. Am. J. Hum. Genet. 103, 338–348. 10.1016/j.ajhg.2018.07.015.

150. Chang, C.C., C.C. Chow, L.C. Tellier, S. Vattikuti, S.M. Purcell, and J.J. Lee (2015). Second-generation PLINK: rising to the challenge of larger and richer datasets. Gigascience 4, 7. 10.1186/s13742-015-0047-8.

151. Picard, M., M.P. Scott-Boyer, A. Bodein, O. Perin, and A. Droit (2021). Integration strategies of multi-omics data for machine learning analysis. Comput Struct Biotechnol J 19, 3735–3746. 10.1016/j.csbj.2021.06.030.

152. Varma, S. and R. Simon (2006). Bias in error estimation when using cross-validation for model selection. BMC Bioinformatics 7, 91. 10.1186/1471-2105-7-91.

153. Campanioni, S., L. Busto, J.A. Gonzalez-Novoa, C. Martinez, P. Juan-Salvadores, I. Vieitez, D.N. Olivieri, J.M. Prieto, I. Vilarino, R. Gonzalez Novas, et al. (2026). Explainable machine learning with bayesian hyper-optimization for predicting cognitive impairment from longitudinal nursing home data. Sci. Rep. 16, 5406. 10.1038/s41598-025-34060-w.

154. Tian, X., G.J. He, P. Hu, L. Chen, C. Tao, Y.L. Cui, L. Shen, W. Ke, H. Xu, Y. Zhao, et al. (2018). Cryptococcus neoformans sexual reproduction is controlled by a quorum sensing peptide. Nat Microbiol 3, 698–707. 10.1038/s41564-018-0160-4.

155. Rathod, R., B. Gajera, K. Nazir, J. Wallenius, and V. Velagapudi (2020). Simultaneous Measurement of Tricarboxylic Acid Cycle Intermediates in Different Biological Matrices Using Liquid Chromatography-Tandem Mass Spectrometry; Quantitation and Comparison of TCA Cycle Intermediates in Human Serum, Plasma, Kasumi-1 Cell and Murine Liver Tissue. Metabolites 10, 10.3390/metabo10030103.

156. Sakuragi, T., R. Kanai, A. Tsutsumi, H. Narita, E. Onishi, K. Nishino, T. Miyazaki, T. Baba, H. Kosako, A. Nakagawa, et al. (2021). The tertiary structure of the human Xkr8-Basigin complex that scrambles phospholipids at plasma membranes. Nat. Struct. Mol. Biol. 28, 825–834. 10.1038/s41594-021-00665-8.

